# Survey of metaproteomics software tools for functional microbiome analysis

**DOI:** 10.1101/2020.01.07.897561

**Authors:** R. Sajulga, C. Easterly, M. Riffle, B. Mesuere, T. Muth, S. Mehta, P. Kumar, J. Johnson, B. Gruening, H. Schiebenhoefer, C. A. Kolmeder, S. Fuchs, B. L. Nunn, J. Rudney, T. J. Griffin, P. D. Jagtap

## Abstract

To gain a thorough appreciation of microbiome dynamics, researchers characterize the functional role of expressed microbial genes/proteins. This can be accomplished through metaproteomics, which characterizes the protein complement of the microbiome. Several software tools exist for analyzing microbiomes at the functional level by measuring their combined proteome-level response to environmental perturbations. In this survey, we explore the performance of six available tools, so that researchers can make informed decisions regarding software choice based on their research goals.

Tandem mass spectrometry-based proteomic data obtained from dental carie plaque samples grown with and without sucrose in paired biofilm reactors were used as representative data for this evaluation. Microbial peptides from one sample pair were identified by the X! Tandem search algorithm via SearchGUI and subjected to functional analysis using software tools including eggNOG-mapper, MEGAN6, MetaGOmics, MetaProteomeAnalyzer (MPA), ProPHAnE, and Unipept to generate functional annotation through Gene Ontology (GO) terms.

Among these software tools, notable differences in functional annotation were detected after comparing differentially expressed protein functional groups. Based on the generated GO terms of these tools we performed a peptide-level comparison to evaluate the quality of their functional annotations. A BLAST analysis against the Universal Protein Knowledgebase revealed that the sensitivity and specificity of functional annotation differed between tools. For example, eggNOG-mapper mapped to the most number of GO terms, while Unipept generated the most precise GO terms. Based on our evaluation, metaproteomics researchers can choose the software according to their analytical needs and developers can use the resulting feedback to further optimize their algorithms. To make more of these tools accessible via scalable metaproteomics workflows, eggNOG-mapper and Unipept 4.0 were incorporated into the Galaxy platform.

## Introduction

Microbiome research has demonstrated the effect of microbiota on their host and environment (1, 2). To determine the key contributors within complex microbiota, nucleic acid-based metagenomics can identify taxonomies that are prevalent in certain environments and stimuli (3). Metagenomics provides an overview of the complete inventory of genes recovered from microbiome samples. As a gene-centric approach it is naturally static and can therefore not reflect temporal dynamics and functional activities of microbiomes. To gain a more impactful understanding of a microbiome, metaproteomics must be used to determine the actions, or functions, of microbial organisms. Specifically, metaproteomics identifies microbial proteins, which are biological units of function (4, 5). Functional analysis also helps in understanding the mechanism by which microorganisms interact with each other and their immediate environment, thus offering deeper insights beyond mere taxonomic composition and correlation of the microbiome(6). For example, functional analysis can provide information which enzymes are active in particular biological processes and metabolic pathways. Thus, investigating the functional metaproteome aims to give biological relevance to the structure of microbiomes and can help to identify metabolism changes caused by specific perturbations and environmental factors.

Metaproteomics data analysis involves primarily identification of peptides from tandem mass spectrometry (MS/MS) data by matching it against protein sequence databases. The identified peptides are assigned to proteins or protein groups, some of which are unique while others are shared amongst various taxa. These identifications are further assigned to functional groups using various annotation databases (7). Compared to single-organism proteomics, the functional annotation of metaproteomics data is not straightforward because it involves multiple complex layers: identified peptides can be assigned to various sequence-similar proteins originating from multiple organisms. This adds on to the already existing challenge of assigning functional groups because proteins can often be assigned to multiple functional groups.

Over the years, multiple software tools have been developed to assess the functional state of a microbial community based on the predicted functions of proteins identified by MS (8–13). These tools differ in various aspects such as – a) input files used for processing, b) annotation databases used for protein and functional assignment, c) peptide-versus protein-level analysis, generated outputs including functional ontology terms, and e) visual outputs generated for biological interpretation (**Figure 1**). Each functional tool has its advantages, and labs across the world have been using them often based on the criterion of how well a certain tool fits into their bioinformatics workflow. However, to our best knowledge, these functional tools have not been compared on the same biological dataset in any benchmarking study yet.

**Figure 1.**
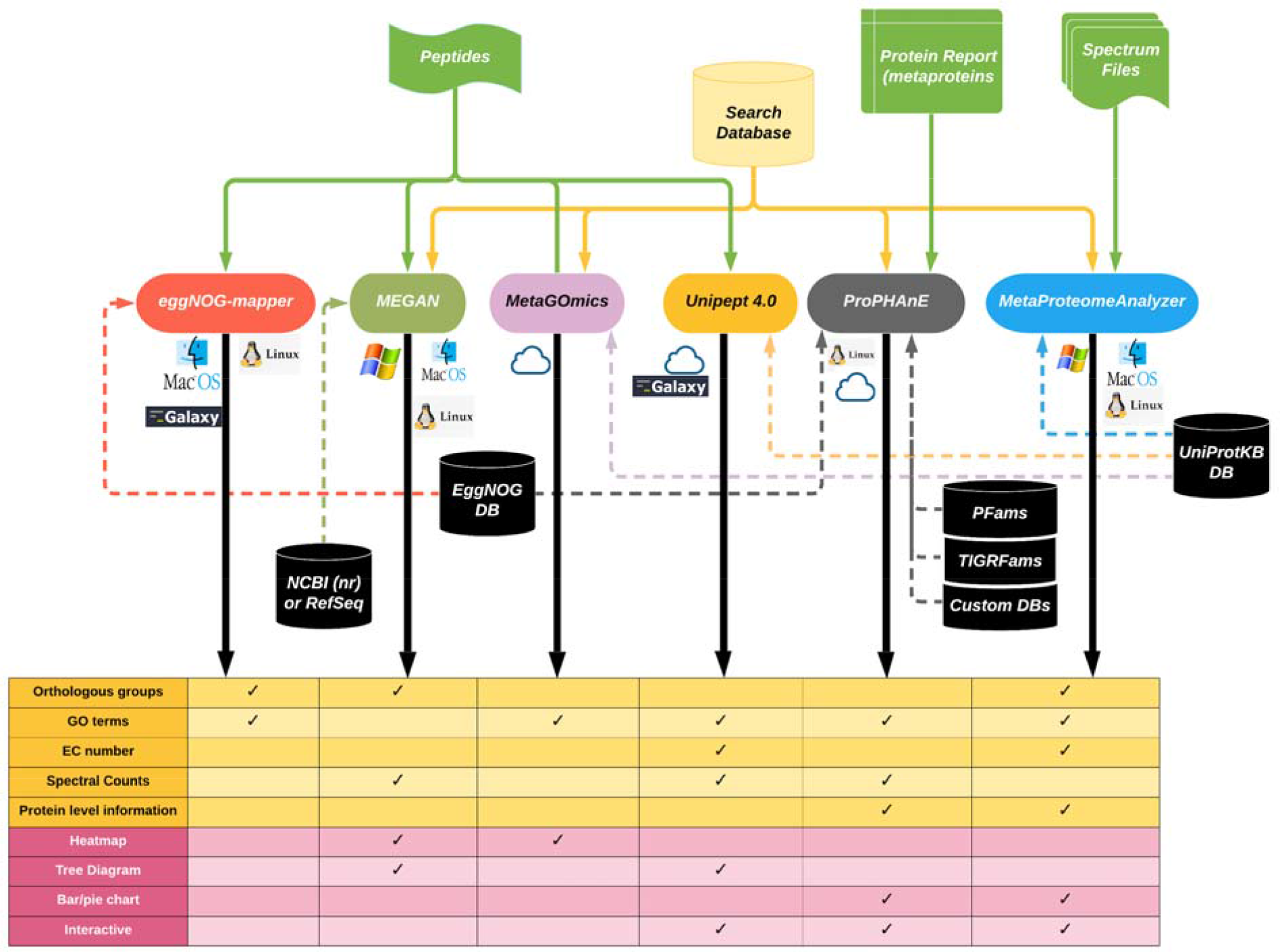
A comparative illustration of the six functional software tools. Required inputs for each tool are connected from the top. Reference databases are connected from the middle. At the bottom, possible output types (data and visualizations) are shown. Next to the tools are labels for supported operating systems and whether the tool has been packaged for Galaxy.

In this study, we compare and evaluate the performance of six open-source software tools - eggNOG-mapper, MEGAN, MetaGOmics, MPA, ProPHAnE and Unipept - that specialize in performing functional analysis of metaproteomics data. For this purpose, we use a published oral dysbiosis dataset(14) to generate outputs and compare features such as identification statistics, functional group assignment (both at a dataset and individual-peptide level), and quantitative analysis features such as differential protein expression. We observed significant variability in results from the different functional tools. Based on further investigation, we provide some insights on the sources of this variability and offer suggestions on the usage of these functional tools.

We have packaged two of these software tools, Unipept 4.0 and eggNOG-mapper, into the Galaxy platform(15), based both on their performance and amenability to deployment in a workflow engine like Galaxy.. This will make these tools widely accessible to users and facilitate their usage in analytical workflows. These software tools are available on GitHub (eggNOG-mapper: https://github.com/galaxyproteomics/tools-galaxyp/tree/master/tools/eggnog_mapper, Unipept: https://github.com/galaxyproteomics/tools-galaxyp/tree/master/tools/unipept), the Galaxy Tool Shed (eggNOG-mapper: https://toolshed.g2.bx.psu.edu/view/galaxyp/eggnog_mapper/, Unipept: https://toolshed.g2.bx.psu.edu/view/galaxyp/unipept) and via Galaxy public instances (https://proteomics.usegalaxy.eu/, http://z.umn.edu/metaproteomicsgateway).

## Methods

The basis for this study lies in selecting a dataset suitable for metaproteomics comparison. Here, we used a published oral dysbiosis dataset (14) (PRIDE PXD003151) containing mass-spectrometry data generated from plaque extracted from dental caries-prone children. These plaques were incubated within biofilm reactors for 72 hours and subject to a pair of conditions for the last 48 hours: with sucrose-pulsing (WS) five times a day and with no sucrose (NS)— only a constant basal mucin medium flow. One sample pair (737 NS/WS) was selected for its quality based on principal component analysis (14). Although not a ‘ground-truth’ dataset, this dataset was chosen since it was already rigorously analyzed (14). From the 737 dataset, MGF (Mascot Generic Format) files containing peak lists were used to identify peptides using X! Tandem search algorithm via SearchGUI. Each of these peptide-spectral matches (PSMs) detail a specific peptide sequence mapped to a specific spectrum scan, as well as match information such as retention time, measured mass-to-charge ratio, theoretical mass and charge, and confidence score. To determine which proteins these identified peptides belong to, a combined protein database was provided: the Human Oral Microbiome Database (HOMD −1,079,742 protein sequences - May 2017) concatenated with the common Repository of Adventitious Proteins (cRAP). Using cRAP proteins to filter out contaminants, HOMD identified ∼27,000 microbial peptides from each dataset (Supplement Data: http://zenodo.org/record/3595577; DOI: 10.5281/zenodo.3595577). Regarding specific SearchGUI parameters, peptides were assigned based on trypsin-digested proteins, with up to two missed cleavage sites allowed. Amino acid modifications were specified as methylthio of cysteine (fixed modification) and oxidation of methionine (variable modification). The accepted precursor mass tolerance was set to 10 ppm and the fragment mass tolerance to 0.05 Da with a charge range from +2 to +6. To interpret SearchGUI results, PeptideShaker was used to generate a PSM, peptide, and protein report, with a false-discovery rate (FDR) cutoff at 1%. Only peptides with length from 6 to 30 amino acids were considered. Spectral counts were calculated based on the number of PSMs assigned for each peptide. Peptide search results were processed to provide appropriate inputs for each of the metaproteomics software tools. (Supplement Data: http://zenodo.org/record/3595577; DOI: 10.5281/zenodo.3595577).

Six functional analysis tools were used to analyze the data: eggNOG-mapper (version 1.0.3), MEGAN5, MetaGOmics (version 0.1.1), MPA (version 1.8.0), Unipept 4.0, and ProPHAnE (3.1.4). Standard procedures were used for each tool, as defined by their developers. Input types are specified in **Figure 1**.

For eggNOG-mapper, we used the Galaxy-implemented version of this tool (v1.0.3) on our local Galaxy for Proteomics (Galaxy-P) server. DIAMOND (Double Index Alignment of Next-generation sequencing Data) was used as a mapping mode since it is much faster, but similar in sensitivity compared to BLAST (Basic Local Alignment Search Tool). PAM30 (Point Accepted Mutation) was used as a scoring matrix (gap costs with an existence value of 9 and extension value of 1). Bacteria were used for taxonomic scope, and all orthologs were considered. Gene ontology evidence was based on non-electronically curated terms, and seed orthology search options had a minimum e-value of 200,000 and a minimum bit score of 20.

For MetaGOmics, a list of peptides and the HOMD were uploaded (version 0.1.1; https://www.yeastrc.org/metagomics). Searches were performed using the UniProt/SwissProt database with a BLAST e-value cutoff at 1e-10, using only the top hit.

In MEGAN, for Lowest Common Ancestor (LCA) analysis we used a minimum score of 30 and a maximum expected threshold value of 3.0. Hits with a BLAST score in the top 10% were chosen for further processing. The minimum support filter to reduce the false positive hits was set as five. Naive LCA algorithm was used with 100% coverage. The read assignment mode was set to read counts and the analysis was performed using eggNOG, InterPro2GO and SEED.

For MPA, the same parameters from SearchGUI: precursor mass tolerance was set at 10 ppm and fragment mass tolerance was set at 0.5 Daltons. Two missed cleavages were allowed and trypsin was used for protein digestion. The false discovery rate (FDR) was set at 1%. For grouping proteins, a minimum of one shared peptide was used if relevant to the analysis.

Unipept takes in a tabular list of unique peptides. Search settings were configured for isoleucine and leucine to be considered distinct, duplicate peptides to be retained, and advanced missed cleavage handling to be enabled.

In ProPHAnE, the NS and WS samples were grouped into one sample group each. Metaproteins (clustered by MPA) were analysed using default parameters. Functional annotation was transferred from TIGRfams v15 and Pfams v32 using HMMScan with trusted cut-off and, additionally, EggNOG v4.5.1 using eggNOG-mapper (hmmer mode).

The six tools described here produce different functional annotation types. To compare and contrast the composition of each tool’s functional annotation, a standard annotation type was chosen. Gene Ontology (GO) terms were used since they are a well-supported and common annotation type throughout most of the tools (i.e., eggNOG-mapper, MetaGOmics, and Unipept) and are commonly used in the metaproteomics community. However, MEGAN, ProPHAnE, and MPA do not provide direct GO term outputs and thus require external databases for translation. MEGAN and ProPHAnE produce eggNOG orthologous group accessions, which are translated using eggNOG’s API (http://eggnogdb.embl.de/#/app/api), while MPA produces UniProt protein accessions, which are used to retrieve UniProtKB GO terms via UniProt’s Retrieve/ID mapping tool (https://www.uniprot.org/uploadlists/).

When using GO terms, it is important to consider their categorization into three different domains: molecular function, biological process, and cellular component. Molecular functions describe the biological activity of gene products at a molecular level (e.g., ATPase activity), biological processes represent widely encompassing pathways that can involve many proteins that aim to accomplish specific biological objectives (e.g., regulation of ATPase activity), and cellular components describe the localization of the activity of these gene products (e.g., plasma membrane). To achieve a high-resolution analysis on functional annotation, only molecular function GO terms were used in this analysis. However, it is important to consider how each tool handles the other ontologies as well.

As an initial evaluation, the GO term outputs of each tool were compared via R scripts (https://github.com/jraysajulga/functional-analysis-benchmarking/blob/master/go_term_compare.Rmd) that looked at the total number of GO terms and the total number of unique terms for each tool. Next, the degree of dissimilarity between the GO term output sets was gauged using fractional overlap indices calculated for each tool pairing. These values were calculated for a tool by taking the size of its intersection set with another tool and dividing by the size of the original tool’s term set. For both of these analyses, GO terms were translated into higher-level, GO generic slim terms to determine if any differences between GO term sets from each tool were due to varying levels of specificity. We used the OBO (Open Biomedical Ontologies) file at http://current.geneontology.org/ontology/subsets/goslim_generic.obo. For reference, there are 147 high-level terms in GO generic slim and 44,945 corresponding terms in GO generic (as of July 1st, 2019).

After comparing the GO term outputs from each tool, fold changes for each term were evaluated (see below). This type of comparison is important in the analysis of microbiomes, since this can reveal which biological functions show an increase or decrease in abundance in response to a stimulus. For our analysis, peptide spectral counts associated with each GO term for both WS and NS conditions were used to estimate fold changes. These counts were available in the peptide reports from PeptideShaker. To ensure fair comparison, spectral count normalization between the two conditions was performed ad hoc. For eggNOG-mapper, MEGAN, and Unipept, a PSM fraction was used to scale down the condition with more spectral counts (NS) to match the other condition (WS): 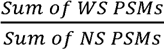. For MPA, we used protein-based normalization, wherein a protein fraction was used to normalize the GO term spectral counts before calculating the log ratio: 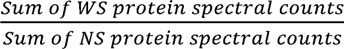. GO terms with multiple instances had their spectral counts aggregated. MetaGOmics and ProPHAnE already internally normalize their values through a normalized spectral abundance factor (NSAF) normalization method (16). Once normalized, these quantitative values were used to calculate the fold changes as 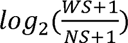, except for MetaGOmics, which already calculated its fold change under the “Log(2) fold change” column name for the comparison output. Using these fold changes, the GO terms were sorted, ascendingly. Thus, GO terms that were found to be more abundant in WS conditions were found at the top of these rankings. Two levels of comparisons of these rankings were used: (1) between tools that natively output GO terms (i.e., Unipept and MetaGOmics); and (2) between all tools, which includes translated GO terms. Initially, Unipept and MetaGOmics were compared by taking the GO terms common between them and plotting each term based on the fold change calculations for Unipept (x-coordinate) and MetaGOmics (y-coordinate). A linear model was regressed, and the Pearson correlation coefficient was calculated with a two-sided alternative hypothesis.

Secondly, all tools were included in a ranking comparison by utilizing translated GO terms. Using the functional annotation outputs from each tool, fold changes were estimated for each molecular function GO term.

With an overall sense of tool discrepancies through identification and quantitation comparisons, the underlying differences between tools were closely examined through a single-peptide analysis. Twenty peptides were randomly selected for this analysis. For a single-peptide analysis, only tools that provided peptide-level outputs were used, thus excluding MEGAN, MPA, and ProPHAnE. The UniProtKB BLAST web search service was used to retrieve GO terms from a peptide (configured with a UniProtKB target database, an expected value threshold of ten, an automatic matrix, no filtering, gap inclusion, and a limit of 50 hits). Sensitivity and specificity of functional annotation differed between tools, in that some tools contained more GO terms than those found through the Universal Protein Knowledgebase (UniProtKB). BLAST, which serves as our standard due to its expert curation, comprehensiveness, and accuracy. Of the peptide-level tools compared against BLAST, Unipept was most similar. Thus, eggNOG-mapper and MetaGOmics, which generally contained many more terms than BLAST/Unipept were scrutinized. To account for these extra terms, the hierarchical structure of GO terms was considered. GO terms assume a top-down hierarchical structure (directed acyclic graphs) which can be navigated using terms such as child, parent, ancestor and descendant. For any GO term, its ancestors consist of any number of less-specialized terms in its hierarchy. Conversely, its descendants consist of more-specific terms that would continue beneath its hierarchy. Parents and children are direct ancestors and direct descendants, respectively (17). As an example, we can examine the ancestors and descendants of ion binding (Go: 0043167): *metal ion binding* (descendant) → *cation binding*(child) → ion binding → binding (parent) → molecular function (ancestor).

To retrieve the ancestors and descendants of GO terms, the Python library GOATOOLS was used (18). Any ancestors, descendants, and ancestors’ children were identified in the extra GO terms found in eggNOG-mapper and MetaGOmics for each peptide. Terms that were uncategorized were labeled as ‘extraneous’.

## Results

### General characteristics of the six tested software

#### Input

The six software tools that were evaluated in this study have been used for metaproteomics analysis before (**Figure 1**). The inputs for each of these tools are different. eggNOG-mapper and Unipept take in only a peptide list and use established databases for annotation (eggNOG and UniProtKB databases, respectively).

In contrast, search databases are required for MEGAN, MetaGOmics, MPA, and ProPHAnE which take in peptides with BLAST-P results, peptides with spectral counts, spectral search files, and spectral search results (from MPA, for example), respectively. MetaGOmics, MPA, and Unipept use UniProtKB as a database for annotation, while MEGAN requires results from NCBI (nr) database. ProPHAnE can use EggNOG, PFAMs, TIGRFams databases and also custom databases for functional annotation.

#### Analysis Level

Moreover, the software tools differ in their level of analysis–some perform analysis at the peptide-level (MetaGOmics, eggNOG-mapper and Unipept) and others perform analysis at the protein or protein-group level (MEGAN, MPA, and ProPHAnE).

#### Outputs

The software tools also generate variable outputs such as proteins annotated with GO terms (eggNOG-mapper, MEGAN, MetaGOmics, MPA, and Unipept 4.0), InterPro2GO (MEGAN), EggNOG orthologous groups (eggNOG-mapper, MEGAN, and ProPHAnE), EC numbers (Unipept 4.0), and Pfam/TIGRFAM accessions (ProPHAnE). Given the variety of inputs, annotation databases, levels of analysis and output types, some variability in results can be expected if the same dataset is processed using these different software tools. To test the degree of variability, we used the published oral dysbiosis microbiome metaproteomics dataset (14).

#### Variation in the number of GO term outputs from functional tools

Peptide reports for the sample grown without sucrose (NS) had 27,420 peptides (56,809 PSMs) and the sample grown with sucrose (WS) had 26,638 peptides (53,205 PSMs). Peptide search results from the oral dysbiosis dataset pair (see methods) were processed to provide appropriate inputs for functional analysis. The outputs from the data processing through these software tools (**Table 2**) show that the total number of functional terms differed for each software tool. To facilitate a fair comparison, the functional terms were standardized into GO terms. The number of total unique GO terms for all ontologies ranged from as low as 1,100 (for MPA) to 6,155 (for eggNOG-mapper). Since this study focuses on functional analysis, we filtered the GO terms to the molecular function ontology. The number of unique molecular function GO terms ranged from as low as 641 (for MPA) to 1,753 (for MetaGOmics). MEGAN and eggNOG-mapper also showed a high number of molecular function GO terms, with eggNOG-mapper exclusively identifying the greatest number of molecular function GO terms (256). The number of slim molecular function GO terms–which offers a higher-level overview of the GO terms–ranged from 31 (MEGAN) to 39 (eggNOG-mapper).

**Table 2:**
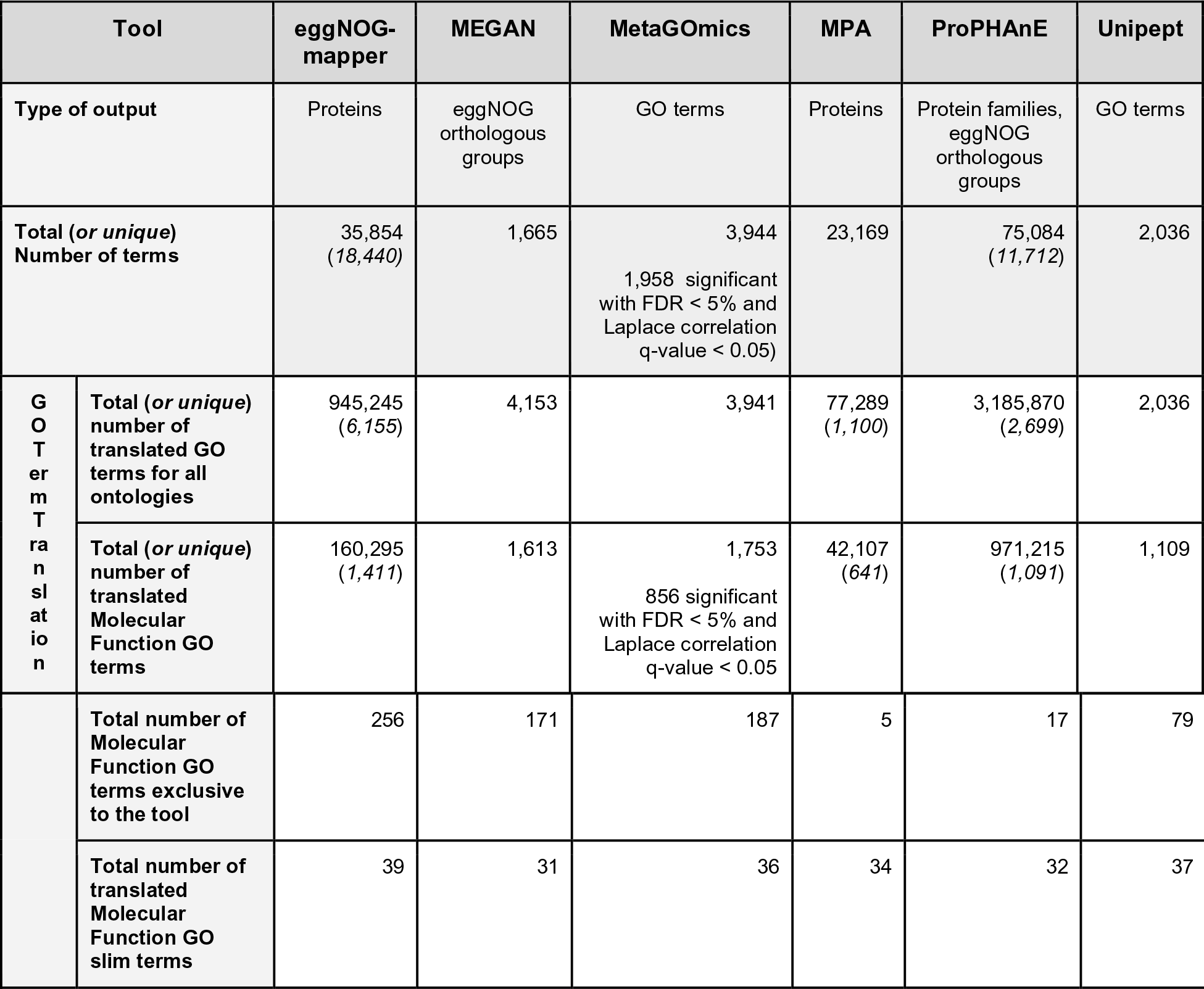
Outputs from functional analysis against the oral dysbiosis dataset

The same analysis was performed for different category combinations as well: all GO terms (molecular function, biological process, and cellular component), biological process GO terms only, and cellular component GO terms only (**<u>Supplement S1)</u>**.

#### Qualitative comparison of functional tools

While comparing the overlap of molecular function GO terms, we found that the tools most similar to each other were MetaGOmics and MPA. MetaGOmics’ GO term set had high coverage in other tools’ GO term sets (with a row-wise average of 0.86) while MPA’s GO term set contained many of the other tools’ terms (with a column-wise average of 0.86). However, MPAs’ GO term set had noticeably lower coverage in other tools (with a row-wise average of 0.41) (**Figure 2A** left panel) due to it having the smallest size (641 terms) as demonstrated in **Table 2**. Relative to GO term comparison, slim GO term comparison shows an improvement due to the more generic (less granular) representation of each term. The same overlap analysis was performed for all GO term ontologies, biological process GO terms, and cellular component GO terms (**<u>Supplement S2</u>**).

**Figure 2:**
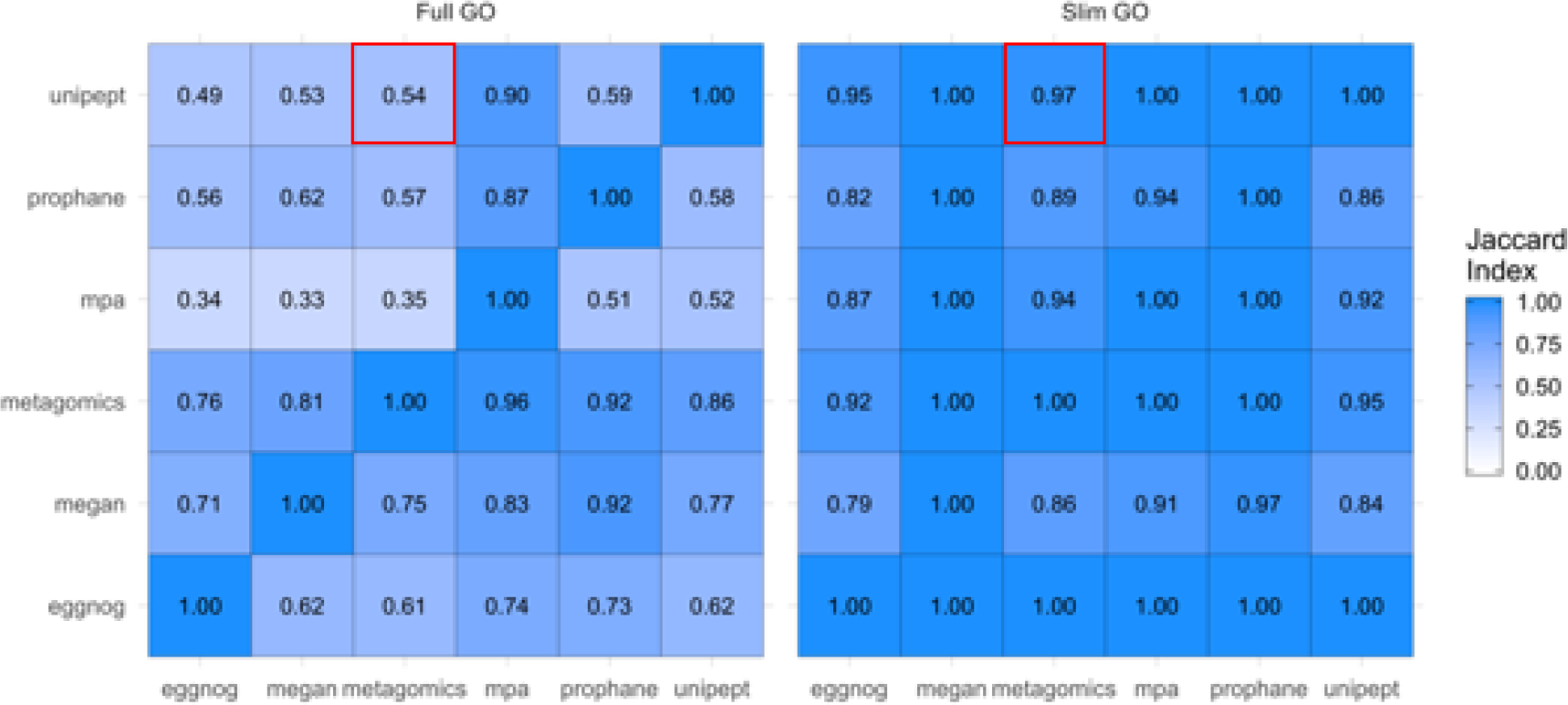
Qualitative and quantitative comparison of functional tools. Overlap of unique GO terms (left) and slim GO terms (right) was compared amongst the six functional tools. Values were calculated as a fraction of the size of term intersection (between the tools labeled on the column and row) over the total term size of the tool listed on the horizontal axis (column). Each functional analysis software tool was compared against each other. For example, for molecular function GO terms (left panel), the fraction of unique Unipept terms present in metaGOmics’ unique GO term set is 0.54 (marked with a red box). For molecular function slim GO terms, the overlap is much larger (0.97).

#### Quantitative comparison of functional tools

After assessing the overlap of functional annotations (**Figure 2A**), we looked at the quantitative changes in GO terms for MetaGOmics and Unipept (**Figure 2B**). As mentioned earlier, MetaGOmics and Unipept both generate GO terms as their output. Comparison of quantitative expression using spectral counts for GO terms from Unipept and MetaGOmics was performed after normalization of spectral counts. Quantitative values of the overlapping molecular function GO terms were represented (**Figure 2B**). The Pearson coefficient of this quantitative comparison was found to be 0.727 with a significant p-value. Given that this is a quantitative comparison of the same dataset, a better quantitative correlation for overlapping molecular function GO terms was expected amongst two functional tools which used the same annotation database (UniProtKB).

**Figure 2 B).**
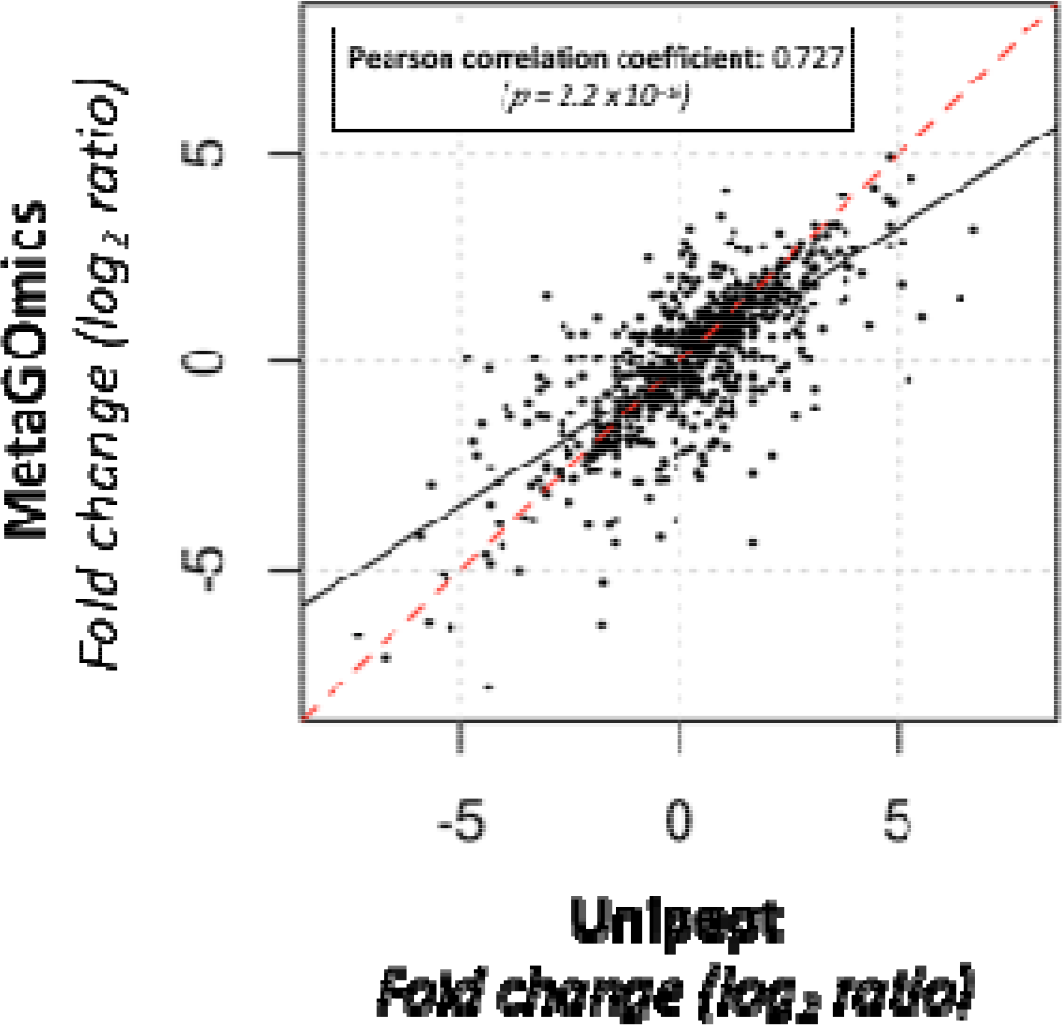
Comparison of quantitative expression for molecular function GO terms from Unipept and MetaGOmics. Log2ratio of spectral counts ‘with sugar sample’ (WS) against ‘no sugar sample’ (NS) was calculated for MetaGOmics- and Unipept-generated molecular function GO terms. Unipept identified 1,109 molecular function GO terms, while MetaGOmics identified 1,753 molecular function GO terms. The data points in figure represent quantitative values for 953 molecular function GO terms that overlapped between Unipept and MetaGOmics.

Quantitative comparison for all GO terms, biological processes GO terms, and cellular component GO terms are available as <u>Supplement S3</u>. The Pearson correlation for all of these comparisons shows lower values than for molecular function category (<u>Supplement S3</u> Figures 1-3).

**Figure 3:**
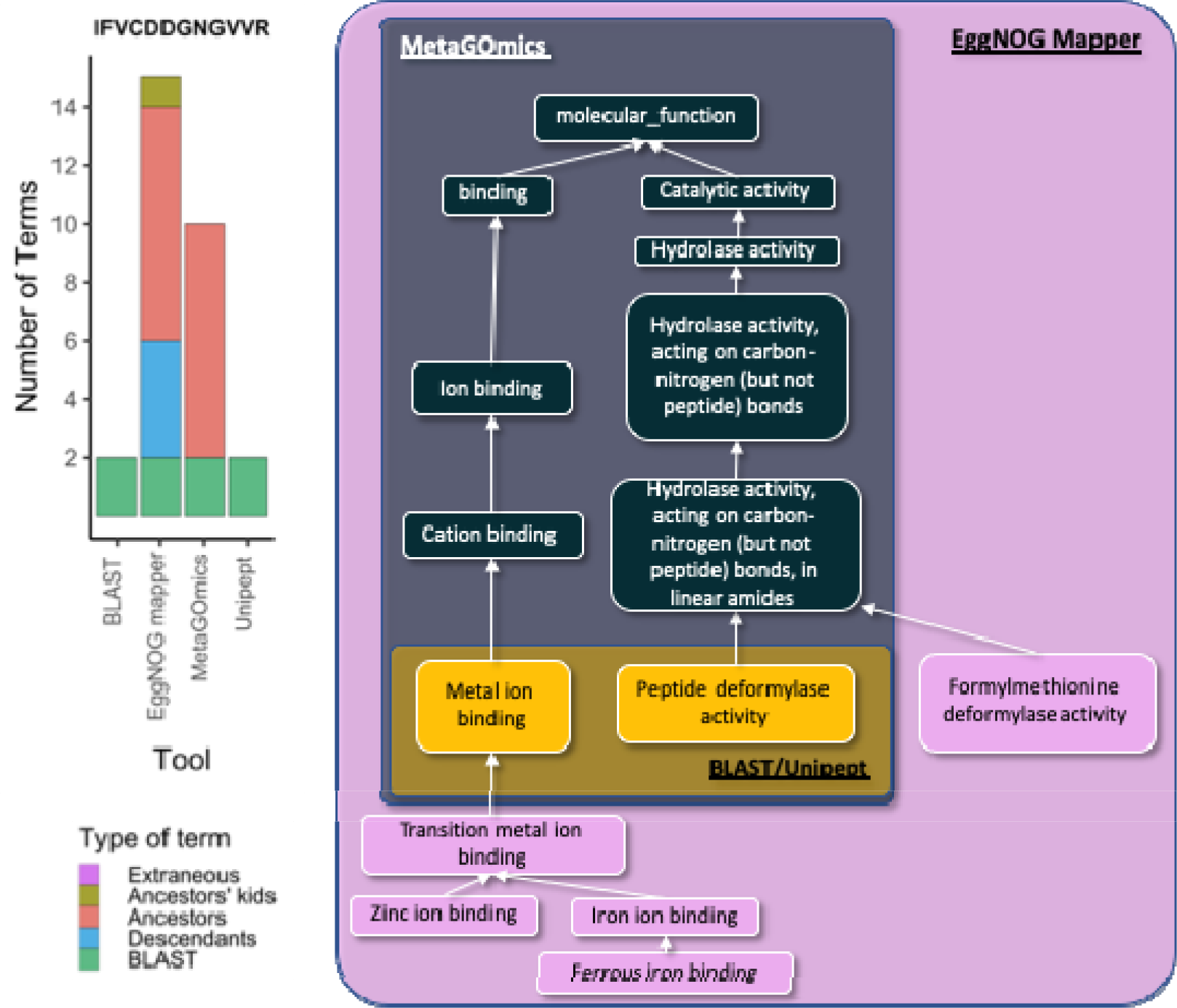
Analysis of peptides associated with the peptide deformylase activity. Functional annotation of a peptide (sequence = IFVCDDGNGVVR) identified in the oral dysbiosis study along with the bar diagram representation of the related terms (descendants, ancestors or ancestor’s children) for eggNOG-mapper, MetaGOmics, Unipept and BLAST-P searches. Molecular function gene ontology hierarchy diagram for a peptide common to BLAST, eggNOG-mapper, MetaGOmics, and Unipept. The terms in the orange box, metal ion binding and peptide deformylase activity, are shared by BLAST/Unipept, MetaGOmics, and eggNOG-mapper. The terms found in the black box are found in both MetaGOmics and eggNOG-mapper. The peripheral terms in the pink box are exclusively annotated by eggNOG-mapper.

**Figure 4:**
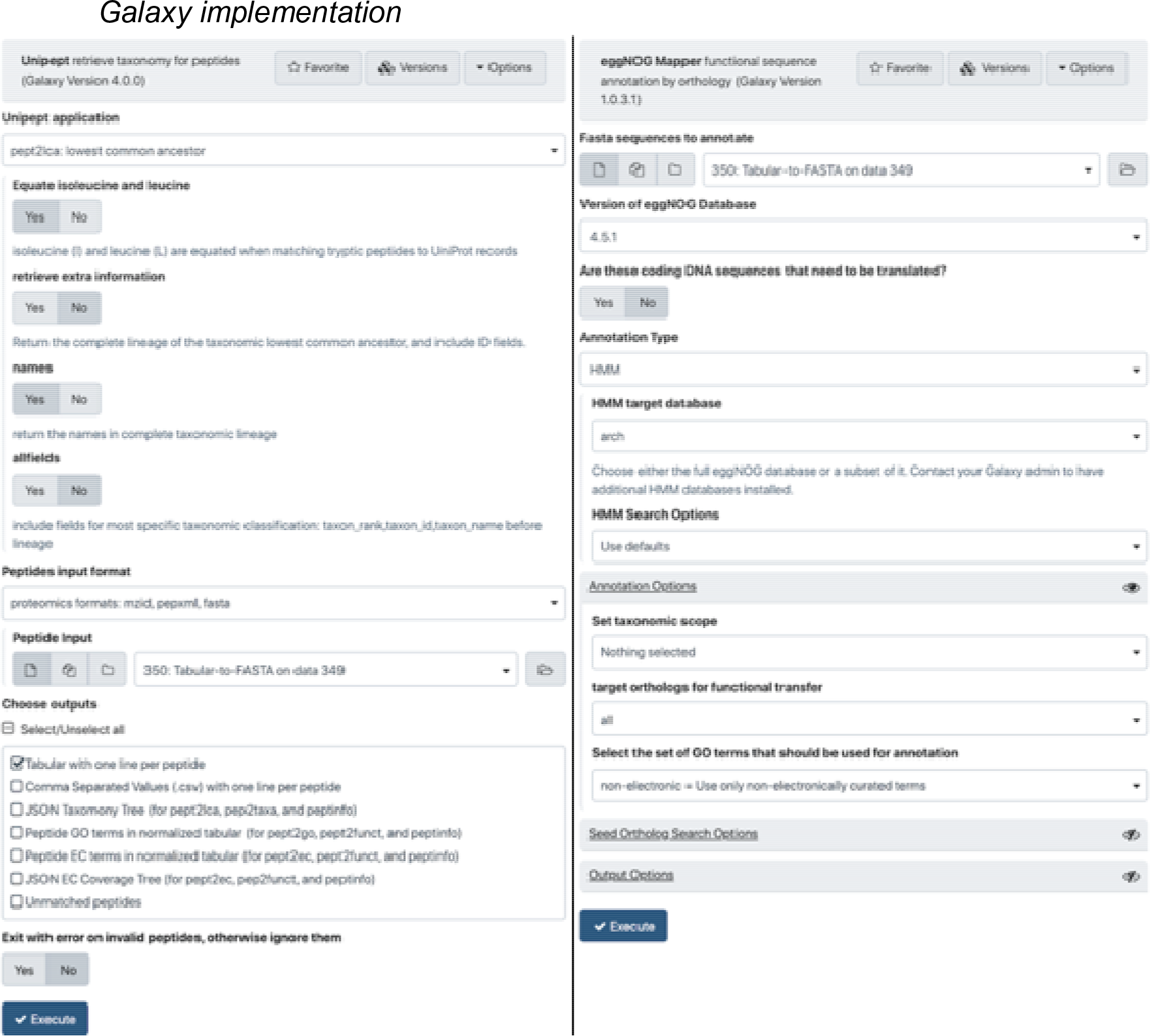
Galaxy interface of Unipept 4.0. Unipept 4.0 is wrapped within Galaxy and available via Galaxy toolshe (https://toolshed.g2.bx.psu.edu/view/galaxyp/unipept), GitHub (https://github.com/galaxyproteomics/tools-galaxyp/tree/master/tools/unipept) and via Galaxy public instances (usegalaxy.eu and z.umn.edu/metaproteomicsgateway).**Galaxy interface of eggNOG-mapper:** eggNOG-mapper is wrapped within Galaxy and available via Galaxy toolshe (https://toolshed.g2.bx.psu.edu/view/galaxyp/eggnog_mapper/), GitHub (https://github.com/galaxyproteomics/tools-galaxyp/tree/master/tools/eggnog_mapper) and Galaxy public instances (usegalaxy.eu and z.umn.edu/metaproteomicsgateway).

A closer look at the fold changes for the molecular function GO terms from the WS and NS data (**Table 3**) revealed that even for the same dataset, when differentially expressed functional groups/terms were ranked, there was substantial variation. While some terms maintained their spectral counts, calculated log fold changes, and ranking statuses for a few functional tools [for example, peptide deformylase activity had 4.93 (#1) and 4.6 (#7) in Unipept and MEGAN, respectively, there were others which varied [for example, 1-phosphofructokinase activity had 4.44 (#2 in Unipept) versus 0.91 (#318 in MPA) fold changes]. This observation was concerning since we expected a better overlap amongst differentially expressed terms amongst functional tools that analyzed the same data (see <u>supplement S4</u> A-E for the rest of the software tools).

**Table 3:**
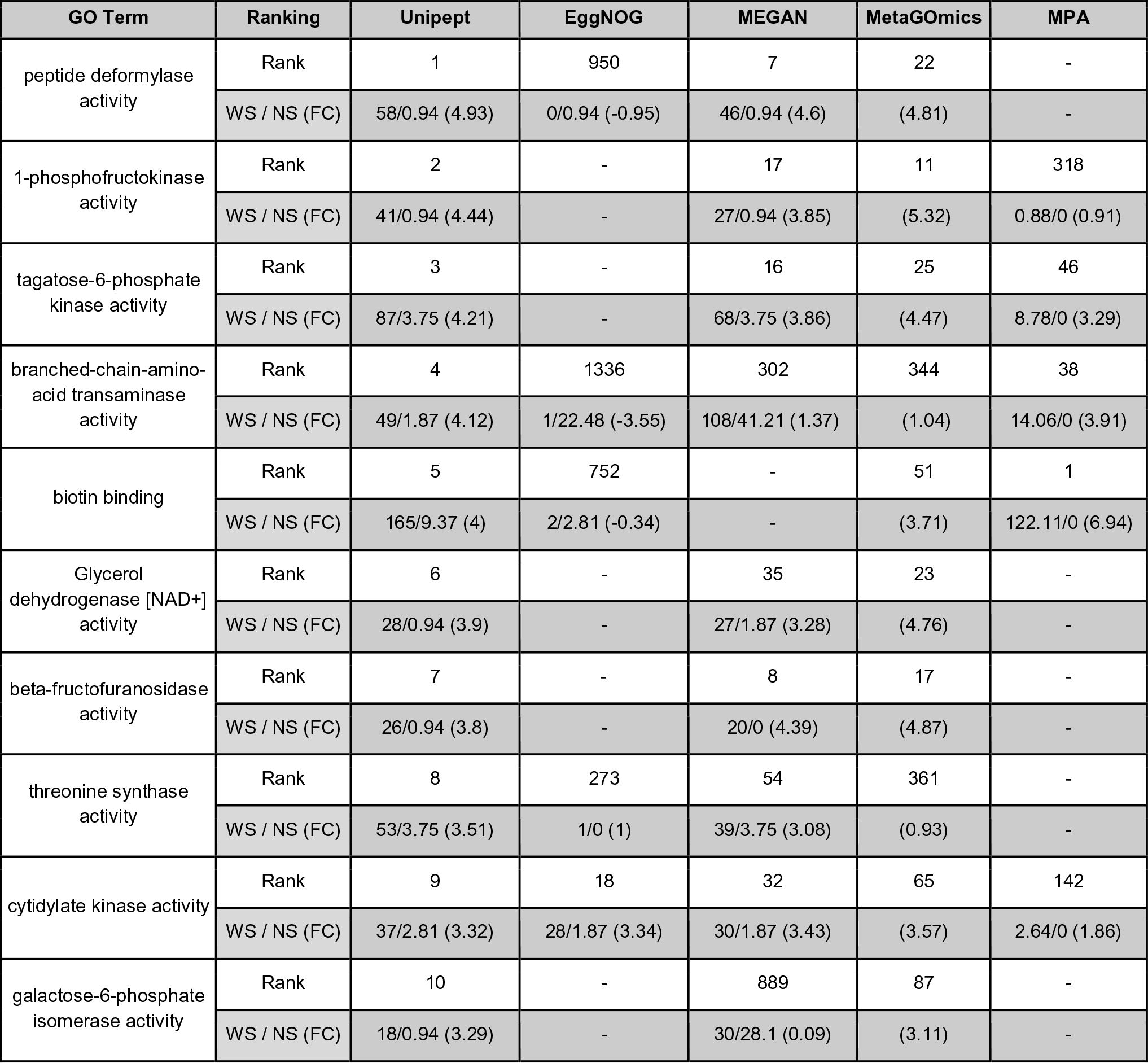
Comparison of the top ten upregulated GO terms of Unipept with the GO terms from the other tools from the oral dysbiosis dataset. The ranks indicate the index of the specific GO term within the list of sorted GO terms based on fold change (descending). Spectral counts are indicated for “with sucrose” and “no sucrose” (WS / NS) conditions which are used to calculate the displayed fold change 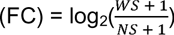. MetaGOmics, however, is compared using its built-in Laplace-corrected fold change value. *ProPHAnE could not identify any of the top ten upregulated molecular function GO terms detected by Unipept. The highest ranks shared between Unipept and ProPHAnE are 14 and 11, respectively, for tagatose-biphosphate aldolase activity. An extended (top 15) version of this table is available in Supplement S4 as Figure S6.*

#### Single peptide analysis

To determine the source of lack of overlap and poor quantitative correlation, we took a deeper look at the functional annotation of the peptide deformylase activity (**Figure 3** and **<u>Supplement</u> <u>S5</u>**) and randomly selected peptides from the oral dysbiosis dataset (**<u>Supplement S6</u>**).

For example, a closer look at the peptides associated with peptide deformylase activity (the top-ranked differentially expressed GO term using Unipept) showed that, while 56 peptides assigned by Unipept, MetaGOmics could assign functional annotation to only 22 peptides (**Table 3**). It should be noted that the PSM ratios from the sucrose-to-control dataset remained similar for Unipept, MetaGOmics and MEGAN. In contrast, eggNOG-mapper could assign only one peptide from the control dataset and could not assign any peptides from the sucrose dataset.

Single peptide analysis of proteins from peptide deformylase activity identified in the oral dysbiosis study (**Figure 3** and <u>Supplement S5</u>) showed that some of the peptides were annotated with related terms (ancestors) by metaGOmics and eggNOG-mapper. Moreover, for most peptides, eggNOG-mapper did not assign any functions. This was in contrast to Unipept and BLAST searches which identified GO terms that were more specific. Bar diagrams for randomly selected peptides from the oral dysbiosis dataset (<u>Supplement S6</u>) also show the number of extraneous, ancestors’ direct children, ancestor terms when using eggNOG-mapper, and MetaGOmics. It is clear that as compared to Unipept (and BLAST), these two functional tools (eggNOG-mapper and metaGOmics) report more functional terms as their outputs. In general, EggNOG contains fewer terms overall compared to MetaGOmics but still contains ancestor terms.

The single-peptide analysis (**Figure 3**, <u>Supplement S5</u> and **<u>Supplement S6</u>**) was carried out for Unipept, MetaGOmics and eggNOG-mapper since they provide GO term annotations at a peptide level. As a result, we do not have results for the protein-level tools MPA and ProPHAnE. However, if peptide-level annotation were enabled for these tools, we anticipate identification of extraneous terms since these software exhibited low Jaccard indices with Unipept (**Figure 2A**) and poor correlation of ranking of differentially expressed proteins (shown for Unipept in **Table 3** and rest of the software tools in Supplement S4).

While assessing the performance of the functional tools presented in this study, we also worked with software developers to explore the possibility of making some of these tools available to metaproteomics researchers within scalable computing environments. To this end, we have implemented software tools and workflows within the Galaxy platform to enable multi-omics analysis (7,19,20). EggNOG-mapper uses orthology assignment, wherein the best matching peptide sequences associated with a protein in the eggNOG database is used to retrieve orthology assignments. In this step, paralogs and matches without sufficient homology are excluded. Subsequently, functions for the retrieved orthologs are transferred to the corresponding query proteins. For eggNOG-mapper implementation within Galaxy, the eggNOG database has to be made accessible along with the interface. The Galaxy data manager application makes it easier for a Galaxy administrator to install reference data. For its usage in Galaxy, users can use DIAMOND against the eggNOG database to generate seed orthologs. The eggNOG-mapper software tool generates outputs such as HMM-hits annotation (if HMM is used as a method) and DIAMOND seed orthologs and annotation (if DIAMOND used as a method).The Galaxy version of eggNOG-mapper is available via Galaxy toolshed (https://toolshed.g2.bx.psu.edu/view/galaxyp/eggnog_mapper/), GitHub (https://github.com/galaxyproteomics/tools-galaxyp/tree/master/tools/eggnog_mapper) and Galaxy public instances (usegalaxy.eu and z.umn.edu/metaproteomicsgateway).

The Unipept software can take in multiple peptide sequences and match them against UniProt database to retrieve taxonomy and functional information. The Unipept Galaxy tool retrieves taxonomy classification and protein-related functional information for peptides using the Unipept API (https://unipept.ugent.be/apidocs). The Unipept tool can read query peptides from a fasta, mzxml, pepxml, or tabular file. The tool has an associated python script that performs the API interaction. (https://github.com/galaxyproteomics/tools-galaxyp/blob/master/tools/unipept/unipept.py) It splits all query peptides into tryptic peptides, queries the Unipept API with the tryptic peptides, then matches the assignments back to the query peptides, generating a tabular format output. Additionally, the output can be a JSON file that can be viewed by the galaxy Unipept tree view visualization. (https://github.com/galaxyproteomics/tools-galaxyp/tree/master/visualizations/unipept) which is an adaptation of the Unipept tree view visualization. (https://github.com/unipept/unipept-visualizations). Within Unipept, a user can perform several kinds of peptide-level analysis including a) access UniProt accession numbers for peptides (pept2prot); b) access taxa information from UniProt entries (pept2taxa and pept2lca); c) access protein EC numbers (pept2ec) and d) access GO term for the peptides (pept2GO). Additionally, Unipept also generates JSON tree outputs that can be used to interactively visualize taxonomic information and EC proteins identified in the sample. Galaxy version of Unipept 4.0 is available via Galaxy toolshed (https://toolshed.g2.bx.psu.edu/view/galaxyp/unipept), GitHub (https://github.com/galaxyproteomics/tools-galaxyp/tree/master/tools/unipept) and via Galaxy public instances (usegalaxy.eu and z.umn.edu/metaproteomicsgateway).

Apart from accessibility, one of the main advantages of having a software tool in Galaxy is the ability to use and evaluate them using repeatable workflows. This allows a user to sequentially connect various tools to generate desired outputs. For example, Unipept and eggNOG-mapper have been used in metaproteomics workflows that can take in MS files and protein database as inputs (21) to eventually generate functional outputs.

## Discussion

Using metaproteomics to investigate microbiomes has gained importance primarily due to its ability to identify functional roles of different taxonomic groups within a complex microbial community. In order to evaluate the software tools that are available for functional analysis of metaproteomics dataset, we used a published dataset of oral dysbiosis from plaque samples derived from dental-carie prone children. In this study, the effect of sucrose on a dental plaque community was assessed wherein principal components analysis showed that the functional content exhibited better separation of the sucrose-treated samples from control samples as compared to taxonomic profiles (14). Although, not a ground truth dataset, we chose this published dataset since it was thoroughly investigated. It is important to note that while taxonomy based ground-truth datasets are available (22), generating a functional microbiome ground truth dataset is not a trivial exercise.

For all of the functional studies, it is important that the software tools used offer results that facilitate a sound biological interpretation. Functional analysis tools either use a peptide-centric (eggNOG-mapper, MEGAN, MetaGOmics and Unipept) or protein-centric approach (MPA and ProPHAnE). Functional annotations are generally performed using functional databases such as Gene Ontology (GO) (17) and Kyoto Encyclopedia of Genes and Genomes (KEGG), or using databases that catalogue the evolutionary relationships of proteins such as orthologous groups (eggNOG). However, all of these databases/approaches are affected by issues associated with annotation quality (4).

Peptides or proteins can also be searched against annotated databases such as NCBI (nr) database (23) and UniProtKB by using algorithms such as BLAST-P or DIAMOND. For taxonomic assignments, peptides unique to a taxonomic unit are used to identify the taxonomic unit(24), while the majority of peptides are assigned at a higher taxonomic level (such as kingdom, phylum, etc.) and cannot be used to identify a lower taxonomic level i.e., strain, species, or genus. In contrast, for functional analysis the identified peptides are assigned to a protein and then a function. This offers an advantage of using functional information of peptides with that have conserved function across taxa, even though they can only be assigned to relatively high taxonomic levels such as kingdom, phylum, etc. However, protein assignments and functional GO term assignments are hierarchical thus making it difficult to assign them to a single function. Moreover, this is confounded by the issue that peptides/proteins can be assigned to multiple functional GO terms.

All of these issues are observed in our results. For example, the database size and composition and underlying algorithm used is reflected in the number of GO terms detected for each software at molecular function and slim GO terms level (**Table 2**). The results from Jaccard index (**Figure 2A**) is also noteworthy, though not surprising considering the variability in database and underlying algorithms used to annotate functions. As expected, Jaccard index improves when using slim GO terms, which have less detailed information. For example, normal GO terms such as “protein disulfide isomerase activity” and “intramolecular oxidoreductase activity” were both translated to the slim GO term “isomerase activity”. We were surprised to see that the quantitative correlation between molecular function terms identified by MetaGOmics and Unipept was not better than 0.72, especially since this correlation was based on the overlapping GO terms. The non-overlap of GO terms (and hence quantitative values) can be explained by the hierarchical nature of GO terms - wherein the same peptides might have been assigned GO terms at varying levels. For overlapping GO terms, the discrepancy in quantitative values might be explained by considering that peptides might have been assigned to different GO terms by different algorithms.

When the ranking of differentially expressed molecular function GO terms was considered, we found a huge discrepancy in the ranking of up-regulated molecular function GO terms (**Table 3**). This also indicated that some of the peptides were assigned to variable GO terms or not assigned at all by some software algorithms. In the case of peptide deformylase activity, which was the top-ranked differentially expressed molecular function GO term using Unipept, we observed that most of the peptides were not assigned using eggNOG-mapper. This may be attributed to the fact that eggNOG-mapper uses fine-grained orthologs to focus on novel sequences. This might have resulted in the loss of quantitative information during eggNOG-mapper analysis for these Unipept-detected ‘Peptide deformylase activity’ peptides and hence the drop in ranking to 132nd position for the differentially expressed gene ontology term. When MetaGOmics was used for analysis of these Unipept assigned peptides, some of the peptides were assigned with ancestor and extraneous terms. The ranking for MetaGOmics analysis of the differentially expressed gene ontology term (‘Peptide deformylase activity’) seems to be in the first decile ranking (rank 11) and not as drastic as eggNOG-mapper (rank 132).

A closer look at the peptides assigned in the study showed the various terms and hierarchy at which molecular function GO terms were assigned (**Figure 3**). This shows the need for filtering of GO terms detected especially when ancestor terms and ancestors’ direct children and extraneous terms were detected. We found that this was also the case for single organism peptides wherein functional annotation depends on matching with a single organism GO term database (data not shown).

Given all of these observations, we noted a few variables that might need to be considered for a more effective functional analysis. There is an opportunity, for example, to improve on the database so that relevant functional information can be easily parsed out. Moreover, methods used to improve on the underlying genome annotation and then transferring the information to the software-specific database is of importance. Next, some of the software seems to generate outputs with additional gene ontology information. This is also augmented by the hierarchical structure of Gene Ontology terms and the issue of multiple GO terms for the same protein. Adding a filtering step after navigating the hierarchical structure of Gene Ontology terms and reporting signal from noise will aid in arriving at fairly consistent outputs.

Based on our knowledge, this is the first study that evaluates the performance of various functional tools in the field of metaproteomics. We acknowledge a few weaknesses in the study, including the basis of this analysis on a single replicate paired-samples and uses spectral counts for quantitative analysis. This design was used primarily to address the current capacities of various software that were compared. Most of the software evaluated in the study support analysis of single-replicate, spectral-count based analysis. We are currently evaluating MS1-based precursor-intensity tools and DIA-based tools for protein quantitation, which we hope will provide more accurate quantitative data for these analysis programs in the future.

While using the same post-search inputs from the same dataset, we were anticipating slight variability in the outputs from the six tools - mainly since they used variable databases for functional annotation along with underlying variability in algorithms. However, we were surprised by the relative less overlap in terms of GO terms (**Figure 2A**), particularly their quantitative outputs (**Figure 2B** and **Table 3**). This was rather noteworthy for tools that generated native GO terms as outputs. A closer look at the peptides and their functional assignment showed that software tools such as Eggnog-mapper and MetaGOmics generated much information that was related but not specific to the protein. Unipept and BLAST, on the other hand showed more focused and specific functional information. As a result of these observations, it is our opinion that in order to measure the functional component of the microbiome, there is a need for further refinement in databases used for annotation and also the underlying algorithms and filtering of the outputs generated from these software tools. This will become increasingly important when evaluating microbiomes from environments where metagenome and metatranscriptome data for peptide spectral matching is available, but not functionally annotated. Metaproteomics researchers face the challenge of analyzing the function of such microbiome systems and software developers have a major role in tackling this challenge. In order to assign functions to such metagenomics data, functional assignment after taxonomic binning is being used. This annotated metagenomics data can be used as a template to assemble metatranscriptomics data and generate protein databases for metaproteomic analysis.

As a result of the study, we have two functional analysis tools which use distinct approaches, database and filtering mechanisms for usage in the Galaxy environment. We hope that the access to these software will encourage metaproteomics researchers to explore these tools either individually or within Galaxy workflows. We also hope that this evaluation encourages software developers to develop tools that generate the right balance of information and filtering so that the functional analysis which is the essence of metaproteomics research can be explored with confidence.

## Acknowledgements

We would like to thank European Galaxy team for the help in the support during Galaxy implementation. We would also like to thank Alessandro Tanca (Porto Conte Ricerche, Italy), Mak Saito and Noelle Held (Woods Hole Oceanographic Institute, Woods Hole, MA) for discussion during the functional tools analysis. We thank Emma Leith for proofreading the manuscript.

We acknowledge funding for this work from the grant National Cancer Institute - Informatics Technology for Cancer Research (NCI-ITCR) grant 1U24CA199347 and National Science Foundation (U.S.) grant 1458524 to T.G. We would also like to acknowledge the Extreme Science and Engineering Discovery Environment (XSEDE) research allocation BIO170096 to P.D.J. and use of the Jetstream cloud-based computing resource for scientific computing (https://jetstream-cloud.org/) maintained at Indiana University. We also acknowledge the support from the Minnesota Supercomputing Institute for maintenance and update of the Galaxy instances. The European Galaxy server that was used for some calculations is in part funded by Collaborative Research Centre 992 Medical Epigenetics (DFG grant SFB 992/1 2012) and German Federal Ministry of Education and Research (BMBF grants 031 A538A/A538C RBC, 031L0101B/031L0101C de.NBI-epi, 031L0106 de.STAIR (de.NBI)).

## Supplemental Data

http://zenodo.org/record/3595577; DOI: 10.5281/zenodo.3595577

Codes: Benchmarking Code and Single Peptide Analysis Code

