## Supplementary Data S1-S6 for "Survey of metaproteomics software tools for functional microbiome analysis"

**Supplement S1**

**Supplement S1 Table 1: All ontologies**

| **Tool** | | **EggNOG mapper** | **MEGAN** | **MetaGOmics** | **MPA** | **ProPHAnE** | **Unipept** |
| --- | --- | --- | --- | --- | --- | --- | --- |
| **Type of annotation** | | Proteins | eggNOG orthologous groups | GO terms | Proteins | Protein families, eggNOG orthologous groups | GO terms |
| **Total number of terms** | | 18,440 | 1,665 | 3,944 (1,958 significant  with FDR < 5% and Laplace correlation q-value < 0.05) | 23,169 | 11,712 | 2,036 |
|  | **Total number of translated GO terms** | 6,155 | 4,153 | 3,941 | 1,100 | 2,699 | 2,036 |
|  | **Number of GO terms exclusive to the tool** | 2,585 | 382 | 407 | 10 | 58 | 175 |
|  | **Total number of translated GO slim terms** | 143 | 129 | 115 | 102 | 110 | 122 |
|  | **Number of GO slim terms exclusive to the tool** | 6 | 0 | 0 | 0 | 0 | 0 |

Peptide search results from the oral dysbiosis dataset pair (see methods) were processed to provide appropriate inputs for functional analysis. The outputs from the data processing through these software tools (Table 2 supplement) shows that the total number of functional terms differed for each software tool. To facilitate a fair comparison, the functional terms were collapsed into GO terms. The number of GO terms for all ontologies ranged from as low as 1100 (for MetaProteomeAnalyzer) to 6155 (for EggNOG mapper).

For all GO terms, the number of Slim GO terms ranged from as less as 102 (for MetaProteomeAnalyzer) to 143 (EggNOG mapper). Notably, EggNOG mapper exclusively identified most number of GO terms (2585) and Slim GO terms (6).

For all GO terms ([**Supplement S1**](https://docs.google.com/document/d/1qR5R7MD9B6wgvQt7s00t-d227UqTSuG_lKKGt9ISLAo/edit) Table 1), the number of slim GO terms ranged from as few as 82 (MEGAN) to as great as 144 (eggNOG-mapper). Notably, eggNOG-mapper exclusively identified the most number of GO terms (3,072) and slim GO terms (nine).

**Supplement S1 Table 2: Biological Process**

| **Tool** | | **EggNOG mapper** | **MEGAN** | **MetaGOmics** | **MPA** | **ProPHAnE** | **Unipept** |
| --- | --- | --- | --- | --- | --- | --- | --- |
| **Type of annotation** | | Proteins | eggNOG orthologous groups | GO terms | Proteins | Protein families, eggNOG orthologous groups | GO terms |
| **Total number of terms** | | 18,440 | 1,665 | 3,944 (1,958 significant  with FDR < 5% and Laplace correlation q-value < 0.05) | 23,169 | 11,712 | 2,036 |
|  | **Total number of translated GO terms (all ontologies)** | 6,155 | 4,153 | 3,941 | 1,100 | 2,699 | 2,036 |
|  | **Total number of translated Biological Processes GO terms** | 3,964 | 2,130 | 1,938  (945 significant  with FDR < 5% and Laplace correlation q-value < 0.05) | 386 | 1,370 | 737 |
|  | **Number of Biological Processes GO terms exclusive to the tool** | 1,928 | 158 | 185 | 5 | 26 | 61 |
|  | **Total number of translated Biological Processes GO slim terms** | 69 | 64 | 55 | 48 | 52 | 57 |
|  | **Number of Biological Processes GO slim terms exclusive to the tool** | 4 | 0 | 0 | 0 | 0 | 0 |

Peptide search results from the oral dysbiosis dataset pair (see methods) were processed to provide appropriate inputs for functional analysis. To facilitate a fair comparison, the functional terms were collapsed into GO terms and were filtered the GO terms to the Biological processes category levels. The number of GO terms for biological process ontology ranged from as low as 386 (for MetaProteomeAnalyzer) to 3964 (for EggNOG mapper). MEGAN and MetaGOmics also showed a high number of molecular function GO terms, with EggNOG mapper exclusively identifying most number of biological process GO terms (1928). The number of Slim biological process GO terms – which offers a higher level overview of the GO terms – ranged from as less as 48 (MetaProteomeAnalyzer) to 69 (EggNOG mapper).

**Supplement S1 Table 3: Cellular component**

| **Tool** | | **EggNOG mapper** | **MEGAN** | **MetaGOmics** | **MPA** | **ProPHAnE** | **Unipept** |
| --- | --- | --- | --- | --- | --- | --- | --- |
| **Type of annotation** | | Proteins | eggNOG orthologous groups | GO terms | Proteins | Protein families, eggNOG orthologous groups | GO terms |
| **Total number of terms** | | 18,440 | 1,665 | 3,944 (1,958 significant  with FDR < 5% and Laplace correlation q-value < 0.05) | 23,169 | 11,712 | 2,036 |
|  | **Total number of translated GO terms (all ontologies)** | 6,155 | 4,153 | 3,941 | 1,100 | 2,699 | 2,036 |
|  | **Total number of translated Cellular Component GO terms** | 640 | 314 | 190  (110 significant  with FDR < 5% and Laplace correlation q-value < 0.05) | 72 | 175 | 178 |
|  | **Number of Cellular Component GO terms exclusive to the tool** | 347 | 45 | 26 | 0 | 7 | 33 |
|  | **Total number of translated Cellular Component GO slim terms** | 34 | 33 | 23 | 19 | 25 | 27 |
|  | **Number of Cellular Component GO slim terms exclusive to the tool** | 1 | 0 | 0 | 0 | 0 | 0 |

Peptide search results from the oral dysbiosis dataset pair (see methods) were processed to provide appropriate inputs for functional analysis. To facilitate a fair comparison, the functional terms were collapsed into GO terms and were filtered the GO terms to the Cellular component category levels. The number of GO terms for biological process ontology ranged from as low as 72 (for MetaProteomeAnalyzer) to 640 (for EggNOG mapper). MEGAN and MetaGOmics also showed a high number of cellular component GO terms, with EggNOG mapper exclusively identifying most number of cellular component GO terms (347). The number of Slim cellular component GO terms ranged from as less as 19 (MetaProteomeAnalyzer) to 34 (EggNOG mapper).

**Supplement S2**

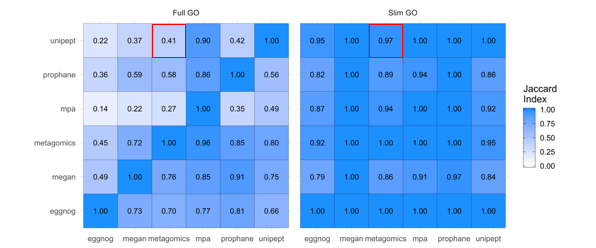

**Supplement 2 Figure 1: Qualitative and quantitative comparison of functional tools (All ontologies)**

A) Overlap of unique GO terms (left) and slim GO terms (right) was compared amongst the six functional tools. Values were calculated as a fraction of the size of term intersection (between the tools labeled on the column and row) over the total term size of the tool listed on the horizontal axis (column). Each functional analysis software tool was compared against each other. For example, for All GO terms (left panel), the fraction of unique Unipept terms present in metaGOmics’ unique GO term set is 0.41 (marked with a red box). For All slim GO terms, the overlap is much larger (0.97).

While comparing the overlap of All GO terms, we found that the tools most similar to other tools were MetaGOmics and MPA. MetaGOmics’ GO term set had high coverage in other tools’ GO term sets (with a row-wise average of 0.76) while MPA’s GO term set contained many of the other tools’ terms (with a column-wise average of 0.87). However, MPAs’ GO term set had noticeably lower coverage in other tools (with a row-wise average of 0.29) (Supplement 2 Figure 1 left panel). Relative to GO term comparison, slim GO term comparison shows an improvement due to the more generic (less granular) representation of each term.

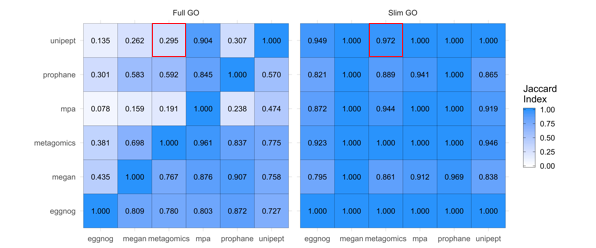

**Supplement 2 Figure 2: Qualitative and quantitative comparison of functional tools (Biological process)**

A) Overlap of unique GO terms (left) and slim GO terms (right) was compared amongst the six functional tools. Values were calculated as a fraction of the size of term intersection (between the tools labeled on the column and row) over the total term size of the tool listed on the horizontal axis (column). Each functional analysis software tool was compared against each other. For example, for biological process GO terms (left panel), the fraction of unique Unipept terms present in metaGOmics’ unique GO term set is 0.295 (marked with a red box). For biological process slim GO terms, the overlap is much larger (0.972).

While comparing the overlap of biological process GO terms, we found that the tools most similar to other tools were MetaGOmics and MPA. MetaGOmics’ GO term set had high coverage in other tools’ GO term sets (with a row-wise average of 0.730) while MPA’s GO term set contained many of the other tools’ terms (with a column-wise average of 0.878). However, MPAs’ GO term set had noticeably lower coverage in other tools (with a row-wise average of 0.228) (Supplement 2 Figure 2 left panel). Relative to GO term comparison, slim GO term comparison shows an improvement due to the more generic (less granular) representation of each term.

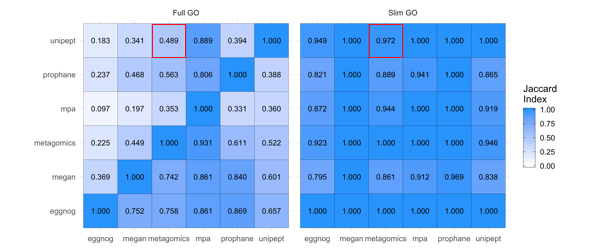

**Supplement 2 Figure 3: Qualitative and quantitative comparison of functional tools (Cellular component)**

A) Overlap of unique GO terms (left) and slim GO terms (right) was compared amongst the six functional tools. Values were calculated as a fraction of the size of term intersection (between the tools labeled on the column and row) over the total term size of the tool listed on the horizontal axis (column). Each functional analysis software tool was compared against each other. For example, for cellular component GO terms (left panel), the fraction of unique Unipept terms present in metaGOmics’ unique GO term set is 0.489 (marked with a red box). For cellular component slim GO terms, the overlap is much larger (0.972).

Cellular component terms had less overlap between tools. MetaGOmics’ GO term set had high coverage in other tools’ GO term sets (with a row-wise average of 0.548) while MPA’s GO term set contained many of the other tools’ terms (with a column-wise average of 0.870). However, MPAs’ GO term set had noticeably lower coverage in other tools (with a row-wise average of 0.268) (Supplement 2 Figure 3 left panel). Relative to GO term comparison, slim GO term comparison shows an improvement due to the more generic (less granular) representation of each term.

**Supplement S3**

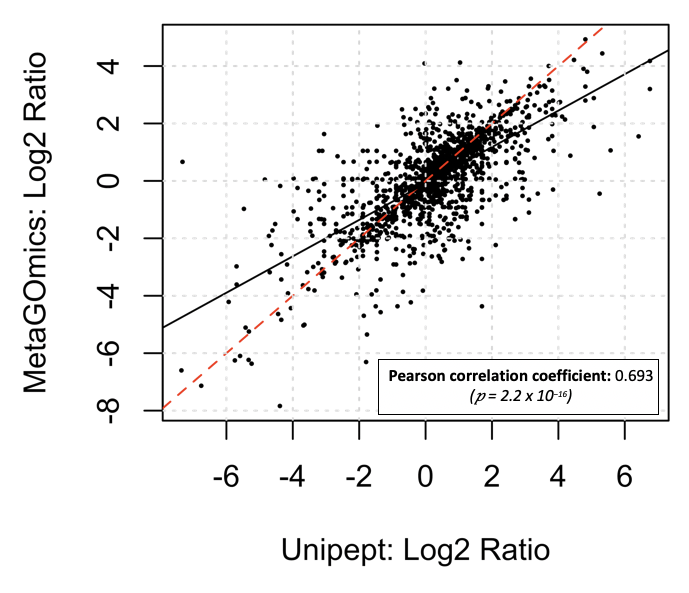

**Supplement S3 Figure 1:** Comparison of quantitative expression for all GO terms from Unipept and MetaGOmics. Log2ratio of spectral counts ‘with sugar sample’ (WS) against ‘no sugar sample’ (NS) was calculated for metaGOmics and Unipept generated all GO terms. Unipept identified 2036 all GO terms, while MetaGOmics identified 3944 all GO terms. The data points in figure represent quantitative values for 1625 all GO terms that overlapped between Unipept and metaGOmics.

Comparison of quantitative expression using spectral counts for all GO terms from Unipept and MetaGOmics was performed after normalization of spectral counts. Quantitative values of the overlapping all GO terms were represented (Supplement S3 Fig 1). The Pearson coefficient of this quantitative comparison was found to be 0.693 with a significant P-value. Given that this is a quantitative comparison of the same dataset, a better quantitative correlation for overlapping all GO terms was expected amongst two functional tools which used the same annotation database (UniProtKB).

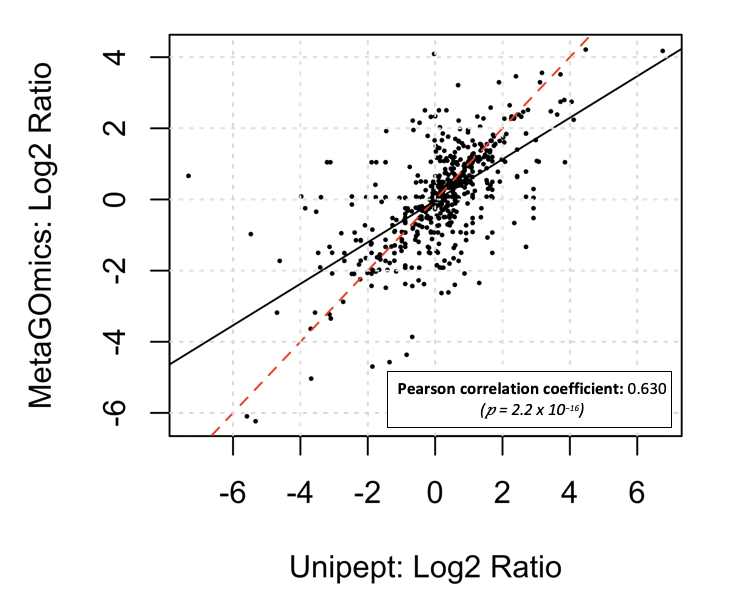

**Supplement S3 Figure 2:** Comparison of quantitative expression for biological process GO terms from Unipept and MetaGOmics. Log2ratio of spectral counts ‘with sugar sample’ (WS) against ‘no sugar sample’ (NS) was calculated for metaGOmics and Unipept generated biological process GO terms. Unipept identified 737 biological process GO terms, while MetaGOmics identified 1938 biological process GO terms. The data points in figure represent quantitative values for 571 biological process GO terms that overlapped between Unipept and metaGOmics.

Comparison of quantitative expression using spectral counts for biological process GO terms from Unipept and MetaGOmics was performed after normalization of spectral counts. Quantitative values of the overlapping biological process GO terms were represented (Supplement S3 Fig 2). The Pearson coefficient of this quantitative comparison was found to be 0.630 with a significant P-value. Given that this is a quantitative comparison of the same dataset, a better quantitative correlation for overlapping biological process GO terms was expected amongst two functional tools which used the same annotation database (UniProtKB).

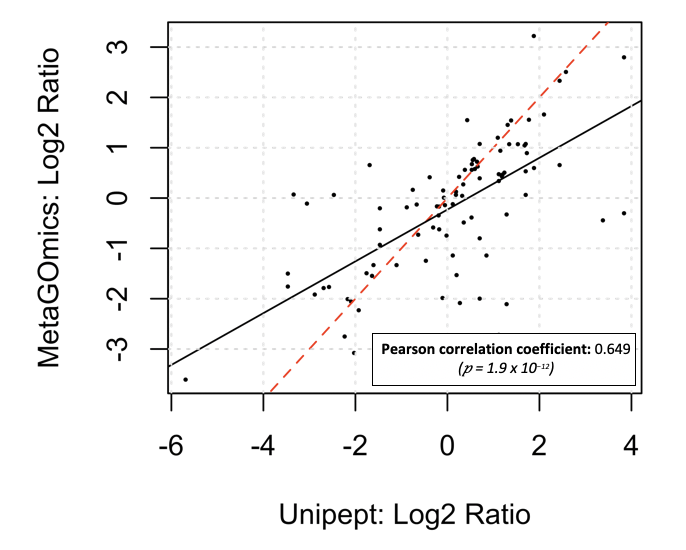

**Supplement S3 Figure 3:** Comparison of quantitative expression for cellular component GO terms from Unipept and MetaGOmics. Log2ratio of spectral counts ‘with sugar sample’ (WS) against ‘no sugar sample’ (NS) was calculated for metaGOmics and Unipept generated cellular component GO terms. Unipept identified 178 cellular component GO terms, while MetaGOmics identified 190 cellular component GO terms. The data points in figure represent quantitative values for 93 cellular component GO terms that overlapped between Unipept and metaGOmics.

Comparison of quantitative expression using spectral counts for cellular component GO terms from Unipept and MetaGOmics was performed after normalization of spectral counts. Quantitative values of the overlapping cellular component GO terms were represented (Supplement S3 Fig 3). The Pearson coefficient of this quantitative comparison was found to be 0.649 with a significant P-value. Given that this is a quantitative comparison of the same dataset, a better quantitative correlation for overlapping cellular component GO terms was expected amongst two functional tools which used the same annotation database (UniProtKB).

**Supplement S4**

**EggNOG**

| **GO Term** | **Ranking** | **EggNOG** | **MEGAN** | **MetaGOmics** | **MPA** | **Prophane** | **Unipept** |
| --- | --- | --- | --- | --- | --- | --- | --- |
| tRNA (5-methylaminomethyl-2-thiouridylate)-methyltransferase activity | Rank | 1 | 224 | - | 759 | - | - |
|  | WS / NS (FC) | 42 / 0 (5.43) | 2 / 0 (1.58) | - | 0 / 3 (-2) | - | - |
| NAD+ synthase activity | Rank | 2 | 533 | 324 | 459 | - | 377 |
|  | WS / NS (FC) | 25 / 0 (4.7) | 21 / 11.24 (0.85) | (1.11) | 4.39 / 4 (0.11) | - | 30 / 18.73 (0.65) |
| aldo-keto reductase (NADP) activity | Rank | 3 | - | - | - | - | - |
|  | WS / NS (FC) | 17 / 0 (4.17) | - | - | - | - | - |
| alcohol dehydrogenase (NADP+) activity | Rank | 4 | - | - | - | - | - |
|  | WS / NS (FC) | 17 / 0 (4.17) | - | - | - | - | - |
| aldehyde dehydrogenase (NADP+) activity | Rank | 5 | 469 | 638 | - | - | - |
|  | WS / NS (FC) | 17 / 0 (4.17) | 1 / 0 (1) | (-1.4) | - | - | - |
| 3-deoxy-7-phosphoheptulonate synthase activity | Rank | 6 | 82 | 111 | - | - | 35 |
|  | WS / NS (FC) | 32 / 0.94 (4.09) | 45 / 6.56 (2.61) | (2.79) | - | - | 55 / 7.49 (2.72) |
| tRNA methyltransferase activity | Rank | 7 | 495 | 310 | - | - | 887 |
|  | WS / NS (FC) | 45 / 1.87 (4) | 57 / 28.1 (1) | (1.19) | - | - | 1 / 2.81 (-0.93) |
| phosphoserine phosphatase activity | Rank | 8 | 176 | - | - | - | - |
|  | WS / NS (FC) | 13 / 0 (3.81) | 16 / 3.75 (1.84) | - | - | - | - |
| phosphatidylserine decarboxylase activity | Rank | 9 | 677 | 641 | - | - | 867 |
|  | WS / NS (FC) | 12 / 0 (3.7) | 3 / 1.87 (0.48) | (-1.44) | - | - | 3 / 6.56 (-0.92) |
| methylglyoxal reductase (NADH-dependent) activity | Rank | 10 | - | - | - | - | - |
|  | WS / NS (FC) | 11 / 0 (3.58) | - | - | - | - | - |

**Supplement S4 Figure 1** Comparison of the top ten upregulated GO terms of eggNOG-mapper with the GO terms from the other tools from the oral dysbiosis dataset. The ranks indicate the index of the specific GO term within the list of sorted GO terms based on fold change (descending). Spectral counts are indicated for “with sucrose” and “no sucrose” (WS / NS) conditions which are used to calculate the displayed fold change (FC) = log_2_($\frac{WS + 1}{NS + 1}$). MetaGOmics, however, is compared using its built-in Laplace-corrected fold change value.

A closer look at the log fold changes for molecular function GO terms from the WS and NS data (S4 Fig 1) revealed that even for the same dataset there was substantial variation between functional tools. Most of the top-ranking terms from eggNOG mapper had a ranking status that was lower in other functional tools. For example, tRNA methyltransferase activity was calculated to have a log fold change of 4.00 (rank #7) while the other tools had lower log fold changes (1.00, 1.19, and -0.93) and much lower ranks (495, 310, 887) for MEGAN, metaGOmics, and Unipept, respectively.

**MEGAN**

| **GO Term** | **Ranking** | **MEGAN** | **EggNOG** | **MetaGOmics** | **MPA** | **ProPHAnE** | **Unipept** |
| --- | --- | --- | --- | --- | --- | --- | --- |
| 2-isopropylmalate synthase activity | Rank | 1 | - | 12 | - | - | - |
|  | WS / NS (FC) | 30 / 0 (4.95) | - | (5.28) | - | - | - |
| homocitrate synthase activity | Rank | 2 | - | - | - | - | - |
|  | WS / NS (FC) | 30 / 0 (4.95) | - | - | - | - | - |
| hydroxymethylglutaryl-CoA lyase activity | Rank | 3 | 149 | - | - | - | - |
|  | WS / NS (FC) | 30 / 0 (4.95) | 2 / 0 (1.58) | - | - | - | - |
| 4-hydroxy-2-oxovalerate aldolase activity | Rank | 4 | - | - | - | - | - |
|  | WS / NS (FC) | 30 / 0 (4.95) | - | - | - | - | - |
| acetolactate synthase activity | Rank | 5 | 869 | 275 | 898 | - | 100 |
|  | WS / NS (FC) | 111 / 2.81 (4.88) | 0 / 0.94 (-0.95) | (1.31) | 0 / 13 (-3.81) | - | 30 / 7.49 (1.87) |
| cysteine-type endopeptidase activity | Rank | 6 | 267 | 389 | 396 | - | 177 |
|  | WS / NS (FC) | 78 / 1.87 (4.78) | 1 / 0 (1) | (0.77) | 29.87 / 21 (0.49) | - | 165 / 60.88 (1.42) |
| peptide deformylase activity | Rank | 7 | 950 | 22 | - | - | 1 |
|  | WS / NS (FC) | 46 / 0.94 (4.6) | 0 / 0.94 (-0.95) | (4.81) | - | - | 58 / 0.94 (4.93) |
| beta-fructofuranosidase activity | Rank | 8 | - | 17 | - | - | 7 |
|  | WS / NS (FC) | 20 / 0 (4.39) | - | (4.87) | - | - | 26 / 0.94 (3.8) |
| sucrose alpha-glucosidase activity | Rank | 9 | - | 18 | - | - | - |
|  | WS / NS (FC) | 20 / 0 (4.39) | - | (4.87) | - | - | - |
| levanase activity | Rank | 10 | - | - | - | - | - |
|  | WS / NS (FC) | 20 / 0 (4.39) | - | - | - | - | - |

**Supplement S4 Figure 2** Comparison of the top ten upregulated GO terms of MEGAN with the GO terms from the other tools from the oral dysbiosis dataset. The ranks indicate the index of the specific GO term within the list of sorted GO terms based on fold change (descending). Spectral counts are indicated for “with sucrose” and “no sucrose” (WS / NS) conditions which are used to calculate the displayed fold change (FC) = log_2_($\frac{WS + 1}{NS + 1}$). MetaGOmics, however, is compared using its built-in Laplace-corrected fold change value.

Most of the top-ranking terms from MEGAN had high log fold changes which were more similar to metaGOmics and Unipept than the other tools (S4 Fig 2). For example, peptide deformylase activity was calculated to have a log fold change of 4.6 (rank #7) for MEGAN while MetaGOmics and Unipept were 4.81 and 4.93, respectively. The same pattern is also seen in beta-fructofuranosidase activity (Rank #8). Other tools did not really detect peptide deformylase activity, except for eggNOG-mapper, which calculated a -0.95 fold change (Rank #950).

**MetaGOmics**

| **GO Term** | **Ranking** | **MetaGOmics** | **EggNOG** | **MEGAN** | **MPA** | **Prophane** | **Unipept** |
| --- | --- | --- | --- | --- | --- | --- | --- |
| pyruvate oxidase activity | Rank | 1 | - | - | - | - | - |
|  | WS / NS (FC) | (7.23 ~ 7.12) | - | - | - | - | - |
| oxidoreductase activity, acting on the aldehyde or oxo group of donors, oxygen as acceptor | Rank | 2 | - | - | - | - | - |
|  | WS / NS (FC) | (7.23 ~ 7.12) | - | - | - | - | - |
| serine-type endopeptidase inhibitor activity | Rank | 3 | 882 | 1220 | 200 | - | 16 |
|  | WS / NS (FC) | (6.76 ~ 6.64) | 0 / 0.94 (-0.95) | 0 / 0.94 (-0.95) | 1.76 / 0 (1.46) | - | 34 / 2.81 (3.2) |
| L(+)-tartrate dehydratase activity | Rank | 4 | - | 167 | 566 | - | 154 |
|  | WS / NS (FC) | (6.42 ~ 6.30) | - | 406 / 105.83 (1.93) | 4.39 / 7 (-0.57) | - | 172 / 58.07 (1.55) |
| ribitol-5-phosphate 2-dehydrogenase activity | Rank | 5 | - | - | - | - | - |
|  | WS / NS (FC) | (6.36 ~ 6.25) | - | - | - | - | - |
| glyceraldehyde-3-phosphate dehydrogenase (NADP+) (non-phosphorylating) activity | Rank | 6 | - | - | - | - | - |
|  | WS / NS (FC) | (6.18 ~ 6.07) | - | - | - | - | - |
| collagen binding | Rank | 7 | 456 | - | - | - | - |
|  | WS / NS (FC) | (6.02 ~ 5.91) | 3 / 1.87 (0.48) | - | - | - | - |
| fructuronate reductase activity | Rank | 8 | - | 948 | - | - | - |
|  | WS / NS (FC) | (5.73 ~ 5.61) | - | 1 / 0.94 (0.05) | - | - | - |
| hyalurononglucosaminidase activity | Rank | 9 | - | - | - | - | - |
|  | WS / NS (FC) | (5.7 ~ 5.58) | - | - | - | - | - |
| transferase activity, transferring acyl groups, acyl groups converted into alkyl on transfer | Rank | 10 | - | 407 | - | - | 249 |
|  | WS / NS (FC) | (5.57 ~ 5.46) | - | 39 / 18.73 (1.02) | - | - | 5 / 1.87 (1.06) |

**Supplement S4 Figure 3** Comparison of the top ten upregulated GO terms of metaGOmics with the GO terms from the other tools from the oral dysbiosis dataset. The ranks indicate the index of the specific GO term within the list of sorted GO terms based on fold change (descending). Spectral counts are indicated for “with sucrose” and “no sucrose” (WS / NS) conditions which are used to calculate the displayed fold change (FC) = log_2_($\frac{WS + 1}{NS + 1}$). MetaGOmics, however, is compared using its built-in Laplace-corrected fold change value (first value; before ‘~’). Here, the spectral count-computed fold changes (second value; after ‘~’) are displayed for validation.

For MetaGOmics, its top ten upregulated terms showed little similarity to other tools (S4 Fig 3). Generally, these terms were not found in most of the other tools. One exception is serine-type endopeptidase inhibitor activity which has a log fold change of 6.76 (#3) in MetaGOmics and is also found in many of the other tools. For eggNOG-mapper, MEGAN, MetaProteomeAnalyzer, ProPHAnE, and Unipept, the calculated log fold changes (and ranks) were -0.95 (#882), -0.95 (#1220), 1.46 (#200), and 3.2 (#16), respectively. Unipept demonstrated the most similarity in this exception.

**MetaProteomeAnalyzer**

| **GO Term** | **Ranking** | **MPA** | **EggNOG** | **MEGAN** | **Meta-**  **GOmics** | **Prophane** | **Unipept** |
| --- | --- | --- | --- | --- | --- | --- | --- |
| biotin binding | Rank | 1 | 752 | - | 51 | - | 5 |
|  | WS / NS (FC) | 122.11 / 0 (6.94) | 2 / 2.81 (-0.34) | - | (3.71) | - | 165 / 9.37 (4) |
| ferroxidase activity | Rank | 2 | 1199 | 1480 | 104 | 463 | 96 |
|  | WS / NS (FC) | 224.89 / 2 (6.23) | 20 / 95.53 (-2.2) | 17 / 86.16 (-2.28) | (2.87) | 0 / 0 (-1.41) | 310 / 81.48 (1.91) |
| heat shock protein binding | Rank | 3 | 594 | - | 209 | - | 168 |
|  | WS / NS (FC) | 41.29 / 0 (5.4) | 2 / 1.87 (0.06) | - | (1.71) | - | 95 / 33.72 (1.47) |
| orotate phosphoribosyltransferase activity | Rank | 4 | - | 887 | - | - | 451 |
|  | WS / NS (FC) | 36.9 / 0 (5.24) | - | 41 / 38.4 (0.09) | - | - | 40 / 29.03 (0.45) |
| pyrimidine nucleobase biosynthetic process | Rank | 5 | - | - | - | - | - |
|  | WS / NS (FC) | 36.9 / 0 (5.24) | - | - | - | - | - |
| adenine phosphoribosyltransferase activity | Rank | 6 | 696 | 653 | 615 | - | 726 |
|  | WS / NS (FC) | 30.75 / 0 (4.99) | 15 / 15.92 (-0.08) | 54 / 36.53 (0.55) | (-1.07) | - | 26 / 32.78 (-0.32) |
| ribonucleoside-diphosphate reductase complex | Rank | 7 | - | - | - | - | - |
|  | WS / NS (FC) | 30.75 / 0 (4.99) | - | - | - | - | - |
| adenine salvage | Rank | 8 | - | - | - | - | - |
|  | WS / NS (FC) | 30.75 / 0 (4.99) | - | - | - | - | - |
| potassium ion transmembrane transporter activity | Rank | 9 | 725 | 658 | 308 | - | 437 |
|  | WS / NS (FC) | 30.75 / 0 (4.99) | 8 / 9.37 (-0.2) | 33 / 22.48 (0.53) | (1.19) | - | 24 / 16.86 (0.49) |
| oxaloacetate metabolic process | Rank | 10 | - | - | - | - | - |
|  | WS / NS (FC) | 28.99 / 0 (4.91) | - | - | - | - | - |

**Supplement S4 Figure 4** Comparison of the top ten upregulated GO terms of MetaProteomeAnalyzer with the GO terms from the other tools from the oral dysbiosis dataset. The ranks indicate the index of the specific GO term within the list of sorted GO terms based on fold change (descending). Spectral counts are indicated for “with sucrose” and “no sucrose” (WS / NS) conditions which are used to calculate the displayed fold change (FC) = log_2_($\frac{WS + 1}{NS + 1}$). MetaGOmics, however, is compared using its built-in Laplace-corrected fold change value.

For MetaProteomeAnalyzer, its top ten upregulated terms showed little similarity to other tools (S4 Fig 4). For example, ferroxidase activity (Rank #2 in MPA) was ranked #96 (Unipept), #104 (MetaGOmics), #463 (Prophane), #1199 (EggNOG mapper) and #1480 (MEGAN).

**Prophane**

| **GO Term** | **Ranking** | **Prophane** | **EggNOG** | **MEGAN** | **Meta-**  **GOmics** | **MPA** | **Unipept** |
| --- | --- | --- | --- | --- | --- | --- | --- |
| 2,3,4,5-tetrahydropyridine-2,6-dicarboxylate N-succinyltransferase activity | Rank | 1 | 994 | 316 | - | 821 | 989 |
|  | WS / NS (FC) | 0 / 0 (3.02) | 2 / 5.62 (-1.14) | 23 / 8.43 (1.35) | - | 0 / 5 (-2.58) | 1 / 5.62 (-1.73) |
| succinyltransferase activity | Rank | 1 | 994 | 316 | 699 | 639 | 866 |
|  | WS / NS (FC) | 0 / 0 (3.02) | 2 / 5.62 (-1.14) | 23 / 8.43 (1.35) | (-2.15) | 0 / 1 (-1) | 3 / 6.56 (-0.92) |
| N-succinyltransferase activity | Rank | 1 | 994 | 316 | - | 821 | 989 |
|  | WS / NS (FC) | 0 / 0 (3.02) | 2 / 5.62 (-1.14) | 23 / 8.43 (1.35) | - | 0 / 5 (-2.58) | 1 / 5.62 (-1.73) |
| tetrahydrodipicolinate N-acetyltransferase activity | Rank | 4 | - | 317 | 108 | 448 | 25 |
|  | WS / NS (FC) | 0 / 0 (3.02) | - | 23 / 8.43 (1.35) | (2.81) | 16.69 / 15 (0.14) | 27 / 2.81 (2.88) |
| N-acetyltransferase activity | Rank | 4 | 554 | 317 | 108 | 250 | 25 |
|  | WS / NS (FC) | 0 / 0 (3.02) | 65 / 58.07 (0.16) | 23 / 8.43 (1.35) | (2.81) | 148.46 / 62 (1.25) | 27 / 2.81 (2.88) |
| N-acyltransferase activity | Rank | 6 | 673 | 811 | 334 | 130 | 1024 |
|  | WS / NS (FC) | 0 / 0 (2.75) | 67 / 67.43 (-0.01) | 104 / 88.04 (0.24) | (1.05) | 19.33 / 4 (2.02) | 9 / 40.27 (-2.05) |
| glutamate-ammonia ligase activity | Rank | 7 | 1340 | 91 | 128 | 79 | 53 |
|  | WS / NS (FC) | 0 / 0 (2.7) | 0 / 12.18 (-3.72) | 160 / 26.22 (2.56) | (2.52) | 252.12 / 36 (2.77) | 187 / 33.72 (2.44) |
| ammonia ligase activity | Rank | 7 | 116 | 91 | 128 | 79 | 53 |
|  | WS / NS (FC) | 0 / 0 (2.7) | 23 / 5.62 (1.86) | 160 / 26.22 (2.56) | (2.52) | 252.12 / 36 (2.77) | 187 / 33.72 (2.44) |
| acid-ammonia (or amide) ligase activity | Rank | 9 | 203 | 188 | 193 | - | - |
|  | WS / NS (FC) | 0 / 0 (2.7) | 48 / 17.79 (1.38) | 252 / 75.86 (1.72) | (1.84) | - | - |
| fructose-bisphosphate aldolase activity | Rank | 10 | 1227 | 802 | - | 504 | 516 |
|  | WS / NS (FC) | 0 / 0 (2.25) | 8 / 43.08 (-2.29) | 387 / 322.18 (0.26) | - | 227.53 / 249 (-0.13) | 408 / 355.89 (0.2) |

**Supplement S4 Figure 5** Comparison of the top ten upregulated GO terms of ProPHAnE with the GO terms from the other tools from the oral dysbiosis dataset. The ranks indicate the index of the specific GO term within the list of sorted GO terms based on fold change (descending). Spectral counts are indicated for “with sucrose” and “no sucrose” (WS / NS) conditions which are used to calculate the displayed fold change (FC) = log_2_($\frac{WS + 1}{NS + 1}$). MetaGOmics, however, is compared using its built-in Laplace-corrected fold change value.

A closer look at the fold changes for molecular function GO terms from the WS and NS data (S4 Fig 5) revealed that even for the same dataset there was substantial variation between functional tools. Most of the top-ranking terms from Prophane had a ranking status that was lower in other functional tools. For example glutamate-ammonia ligase activity ranked #7 in Prophane is ranked 53rd in Unipept, 79th in MetaProteomeAnalyzer, 91^st^ in MEGAN and #128th in metaGOmics).

**Unipept (Top 15)**

| **GO Term** | **Ranking** | **Unipept** | **EggNOG** | **MEGAN** | **MetaGOmics** | **MPA** | **Prophane** |
| --- | --- | --- | --- | --- | --- | --- | --- |
| peptide deformylase activity | Rank | 1 | 950 | 7 | 22 | - | - |
|  | WS / NS (FC) | 58 / 0.94 (4.93) | 0 / 0.94 (-0.95) | 46 / 0.94 (4.6) | -4.81 | - | - |
| 1-phosphofructokinase activity | Rank | 2 | - | 17 | 11 | 318 | - |
|  | WS / NS (FC) | 41 / 0.94 (4.44) | - | 27 / 0.94 (3.85) | -5.32 | 0.88 / 0 (0.91) | - |
| tagatose-6-phosphate kinase activity | Rank | 3 | - | 16 | 25 | 46 | - |
|  | WS / NS (FC) | 87 / 3.75 (4.21) | - | 68 / 3.75 (3.86) | -4.47 | 8.78 / 0 (3.29) | - |
| branched-chain-amino-acid transaminase activity | Rank | 4 | 1336 | 302 | 344 | 38 | - |
|  | WS / NS (FC) | 49 / 1.87 (4.12) | 1 / 22.48 (-3.55) | 108 / 41.21 (1.37) | -1.04 | 14.06 / 0 (3.91) | - |
| biotin binding | Rank | 5 | 752 | - | 51 | 1 | - |
|  | WS / NS (FC) | 165 / 9.37 (4) | 2 / 2.81 (-0.34) | - | -3.71 | 122.11 / 0 (6.94) | - |
| glycerol dehydrogenase [NAD+] activity | Rank | 6 | - | 35 | 23 | - | - |
|  | WS / NS (FC) | 28 / 0.94 (3.9) | - | 27 / 1.87 (3.28) | -4.76 | - | - |
| beta-fructofuranosidase activity | Rank | 7 | - | 8 | 17 | - | - |
|  | WS / NS (FC) | 26 / 0.94 (3.8) | - | 20 / 0 (4.39) | -4.87 | - | - |
| threonine synthase activity | Rank | 8 | 273 | 54 | 361 | - | - |
|  | WS / NS (FC) | 53 / 3.75 (3.51) | 1 / 0 (1) | 39 / 3.75 (3.08) | -0.93 | - | - |
| cytidylate kinase activity | Rank | 9 | 18 | 32 | 65 | 142 | - |
|  | WS / NS (FC) | 37 / 2.81 (3.32) | 28 / 1.87 (3.34) | 30 / 1.87 (3.43) | -3.57 | 2.64 / 0 (1.86) | - |
| galactose-6-phosphate isomerase activity | Rank | 10 | - | 889 | 87 | - | - |
|  | WS / NS (FC) | 18 / 0.94 (3.29) | - | 30 / 28.1 (0.09) | -3.11 | - | - |
| ADP-ribose diphosphatase activity | Rank | 11 | 960 | 1278 | 72 | - | - |
|  | WS / NS (FC) | 18 / 0.94 (3.29) | 0 / 0.94 (-0.95) | 0 / 0.94 (-0.95) | -3.43 | - | - |
| 6-phosphogluconolactonase activity | Rank | 12 | 36 | 178 | 19 | - | - |
|  | WS / NS (FC) | 27 / 1.87 (3.28) | 7 / 0 (3) | 29 / 7.49 (1.82) | -4.81 | - | - |
| proline dipeptidase activity | Rank | 13 | - | - | - | 221 | - |
|  | WS / NS (FC) | 100 / 9.37 (3.28) | - | - | - | 1.76 / 0 (1.46) | - |
| tagatose-bisphosphate aldolase activity | Rank | 14 | 1169 | 322 | 64 | 53 | 11 |
|  | WS / NS (FC) | 45 / 3.75 (3.28) | 0 / 2.81 (-1.93) | 427 / 172.33 (1.3) | -3.6 | 7.91 / 0 (3.15) | 0 / 0 (2.25) |
| transferase activity, transferring phosphorus-containing groups | Rank | 15 | 523 | 709 | 511 | 166 | 257 |
|  | WS / NS (FC) | 43 / 3.75 (3.21) | 1584 / 1319.61 (0.26) | 5090 / 3781.83 (0.43) | -0.26 | 21.08 / 6 (1.66) | 0.02 / 0.01 (0.49) |

**Supplement S4 Figure 6** Comparison of the top fifteen upregulated GO terms of Unipept with the upregulated GO terms from the other tools from the oral dysbiosis dataset. The ranks indicate the index of the specific GO term within the list of sorted GO terms based on fold change (descending). Spectral counts are indicated for “with sucrose” and “no sucrose” (WS / NS) conditions which are used to calculate the displayed fold change (FC) = log_2_($\frac{WS + 1}{NS + 1}$).

**Supplement S5**

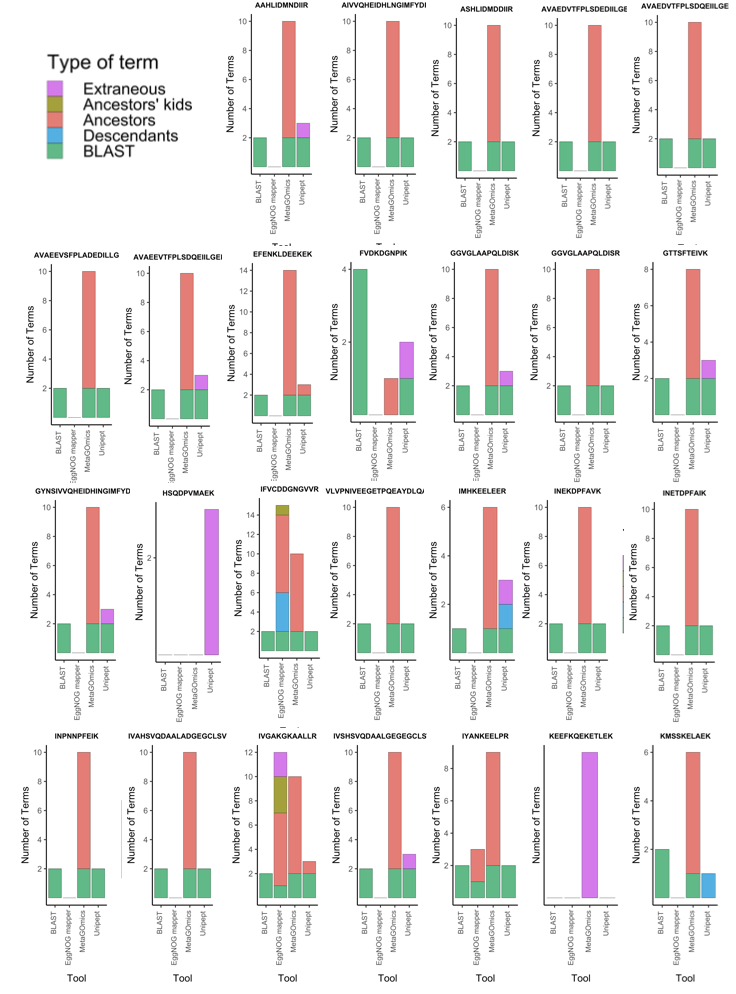

Distribution of molecular function GO terms of eggNOG mapper and metaGOmics for 26 peptides from the oral dysbiosis dataset with molecular function GO term as ‘Peptide deformylase activity’. Molecular function GO terms found in common with BLAST and Unipept peptide results were set as an anchor point at the bottom (green). The rest of the GO terms were hierarchically categorized relative to these BLAST/Unipept-intersected GO terms e.g., descendants (blue) and ancestors (red). Ancestors’ kids are any terms that are the direct descendants of any ancestors of any BLAST/Unipept-intersected GO term in each peptide. Extraneous terms (purple) are any terms that do not fit in any of the aforementioned categories.

**Supplement S6**
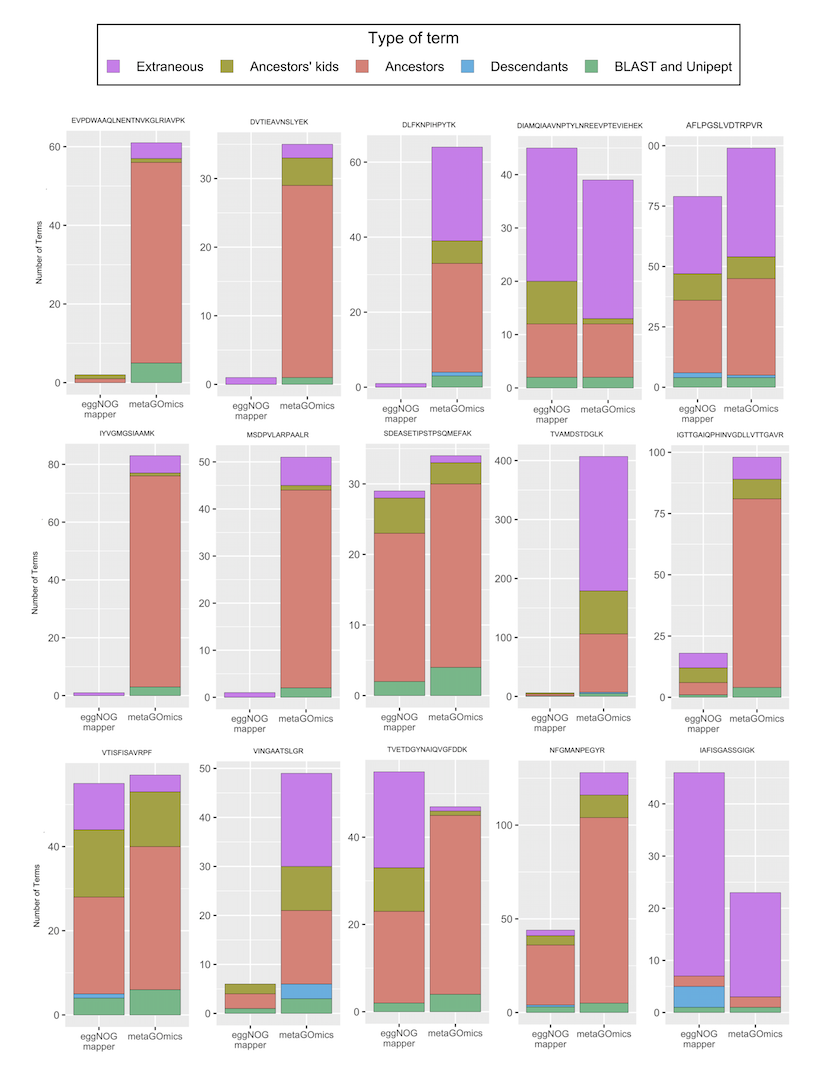

Distribution of molecular function GO terms of eggNOG mapper and metaGOmics for 15 randomly selected peptides from the oral dysbiosis dataset. Molecular function GO terms found in common with BLAST and Unipept peptide results were set as an anchor point at the bottom. The rest of the GO terms were hierarchically categorized relative to these BLAST/Unipept-intersected GO terms e.g., descendants and ancestors. Ancestors’ kids are any terms that are the direct descendants of any ancestors of any BLAST/Unipept-intersected GO term in each peptide. Extraneous terms are any terms that do not fit in any of the aforementioned categories.
